# Time-resolved ligand dynamics revealed in a β-lactamase using room-temperature serial crystallography

**DOI:** 10.64898/2025.12.17.692331

**Authors:** Philip Hinchliffe, Catherine L. Tooke, Michael Beer, Pierre Aller, Jos J. A. G. Kamps, Laura Parkinson, Tiankun Zhou, Nicholas Devenish, Do-Heon Gu, Anastasya Shilova, Matthew J. Rodrigues, Abbey Telfer, Agata Butryn, Marko Hanževački, Emily I. Freeman, Sehan Park, Christopher J. Schofield, Jaehyun Park, Robin L. Owen, Adrian J. Mulholland, Allen M. Orville, James Spencer

**Author notes:** Corresponding authors: Philip Hinchliffe; Allen M. Orville; James Spencer. Equal contribution (Philip Hinchliffe, Allen M. Orville, James Spencer). **Competing interests:** The authors declare no competing interest.

## Abstract

β-Lactamases catalyze β-lactam antibiotic hydrolysis and are important contributors to bacterial antimicrobial resistance; β-lactamase inhibitors are widely used to overcome β-lactamase-mediated antibiotic resistance. Nucleophilic serine β-lactamases (SBLs) react with their substrates and clinically available inhibitors via a covalent reaction to give complexes which can undergo further reaction. Using room temperature drop on fixed target serial crystallography, where ligands are rapidly mixed with microcrystals, and classical single-crystal crystallography at cryogenic temperatures, we investigate the reversible covalent reaction of the SBL CTX-M-15 with the diazobicyclooctane inhibitor avibactam. We observe avibactam covalently reacted (ring-opened) with the nucleophilic Ser70, at timepoints from 80 ms to minutes (room temperature) and hours (100 K). These crystallographic data reveal time-dependent movement of the avibactam carbamoyl complex, from 1.3 s onwards, that has implications for the 5-exo-trig recyclization mechanism that determines inhibitor reformation. Combined with molecular dynamics simulations and quantum mechanics calculations at the density functional theory level, the results show that in the first seconds of the reaction the avibactam *N*-sulfate nitrogen is poorly positioned for recyclization. This subsequently equilibrates after 10 s to a stable endpoint that is in a conformation potentially primed to initiate recyclization through attack of the *N*-sulfate nitrogen on the carbamoyl carbon. These results further demonstrate the capacity of room-temperature serial crystallography to capture time-resolved changes in ligand conformation at an enzyme active site, complementing discrete classical cryo-crystallography. These data inform on ligand dynamics and the stereoelectronics of diazobicyclooctane inhibition, aiding drug discovery efforts to develop inhibitors of nucleophilic enzymes.

## Introduction

Antimicrobial resistance (AMR) is estimated to contribute to 4.95 million global deaths in 2019, with 1.27 million of these directly attributable to drug-resistant infections (1). The production of β-lactamases is the most important AMR determinant in Gram-negative bacteria (2) as these enzymes collectively catalyze the breakdown of all clinically used β-lactams, the most prescribed antibiotic class (3). β-Lactamases (EC 3.5.2.6) are a wide and diverse enzyme family which inactivate β-lactams through hydrolysis (**Figure 1A**). These enzymes are divided into 4 main classes (A-D) based on sequence, structure and mechanism (4). CTX-M-15 is a class A serine β-lactamase (SBL) that hydrolyzes penicillins and oxyimino-cephalosporins (5, 6), which is distributed worldwide on plasmids that are hosted in multiple species of pathogenic Gram-negative bacteria (7). Enterobacterales (*Escherichia coli* and related species) that produce CTX-M-15, and similar so-called extended spectrum β-lactamases (ESBLs), are classed by the World Health Organisation as Critical Priority pathogens for antibacterial research and development.

**Figure 1.**
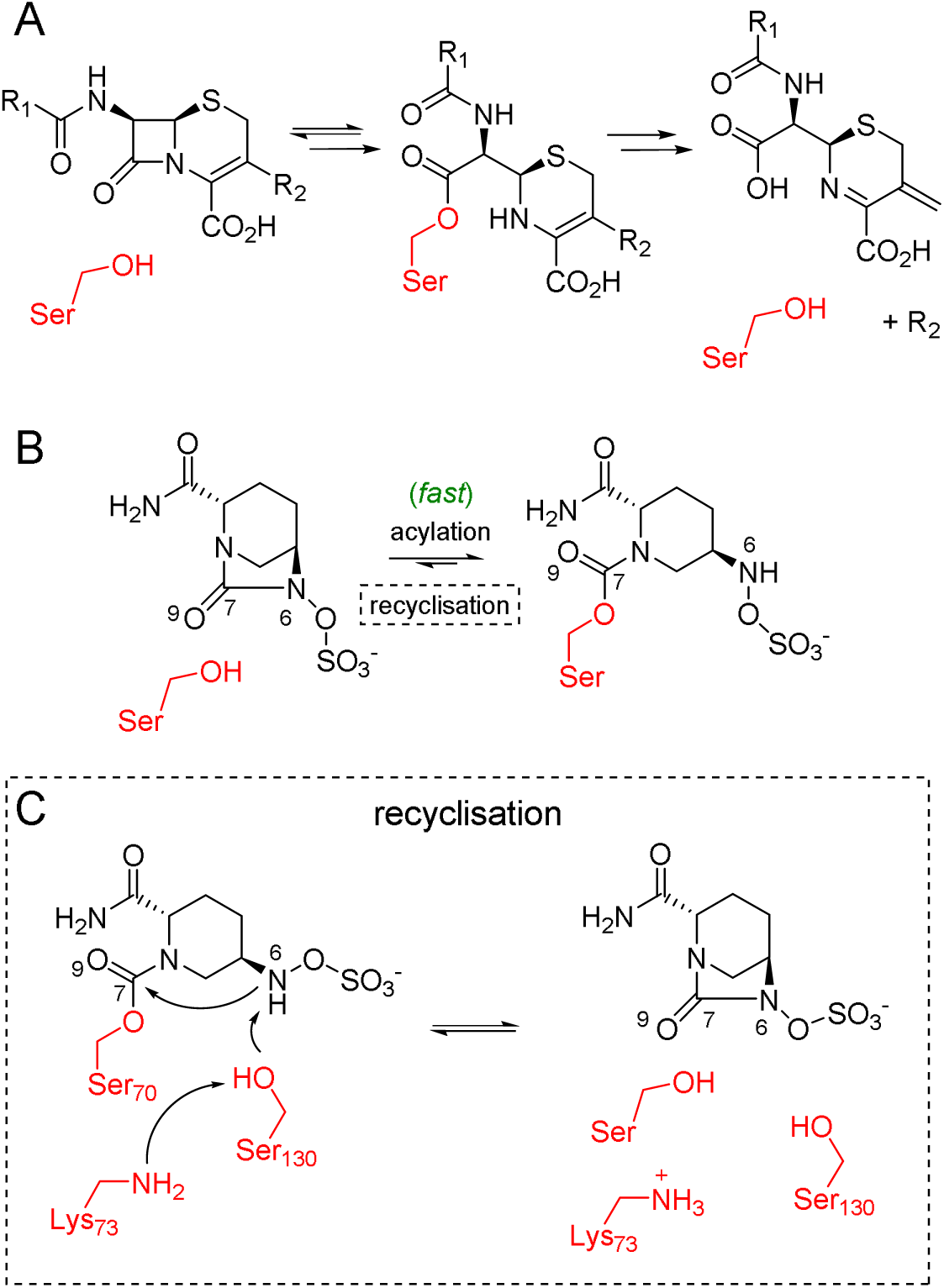
Reactions of β-lactams and avibactam with class A serine-β-lactamases. **(A)** Simplified mechanism of β-lactam hydrolysis and ring-opening by serine-β-lactamases, exemplified by hydrolysis of a cephalosporin antibiotic, via formation of an acyl-enzyme intermediate, resulting in loss of the C3 R_2_ substituent and formation of an exocyclic methylene group. **(B)** Reaction of avibactam with the Ser70 nucleophile, resulting in formation of the carbamoyl enzyme complex, after breakage of the N6-C7 bond. Protonation of N6 likely occurs by Ser130. **(C)** Proposed recyclization mechanism, involving proton transfers between a neutral Lys73 (based on work in (19–21), see Introduction), Ser130 and the avibactam NH (N6) before attack on the C7 carbonyl carbon and reformation of the active site and intact avibactam (right). Note, for optimal N6-C7-O9 burgi-durnitz angle for carbonyl attack is 107°.

To circumvent the action of β-lactamases, inhibitors have been partnered with β-lactams in the clinic to restore bactericidal activity (8). Avibactam, the first approved member of the emerging and expanding diazabicyclooctane (DBO) class of non-β-lactam β-lactamase inhibitors, was introduced in 2015 in combination with the oxyimino-cephalosporin ceftazidime (**Figure S1A**) (9). In 2017, a closely related DBO with a modified C2 amido side chain, relebactam (**Figure S1B**), was introduced, but although efficacious in combination with imipenem-cilastatin against some bacterial infections, it has a lower affinity for class A β-lactamases than avibactam (10). Due to their clinical success there is increasing attention on developing novel DBOs (11), leading to the identification of some with dual action capability that encompasses the penicillin binding proteins, targets for β-lactam action, as well as SBLs. These include nacubactam (12, 13), zidebactam (14), WCK-5153 (15), ETX-0462 (16) and durlobactam (previously ETX-2514 (17)), which have modifications at C2, C3 and C4 of the DBO ring system (**Figure S1C**), and which have demonstrated antimicrobial activity. In 2023, the combination of durlobactam and the β-lactam-based β-lactamase inhibitor sulbactam was introduced to the clinic to treat *Acinetobacter* infections (18). DBOs are thus of increasing importance in the fight against AMR.

Avibactam was a pioneering example of a clinically approved nucleophilic enzyme inhibitor working by a reversible covalent ‘acylation type’ mechanism. It, and other DBOs, inhibit SBLs by forming a stable, covalent carbamoyl ester following attack by the nucleophilic serine residue - Ser70 in CTX-M-15 (**Figure 1B**) (8, 22). The inhibition of SBLs by avibactam has been studied in detail, revealing it as a highly potent inhibitor that rapidly reacts covalently with many SBLs, with an IC_50_ value against CTX-M-15 of 3.4 nM (10), a carbamoylation/on rate of 1.3 ± 0.1 × 10^5^ M^-1^ s^-1^, a *k*_off_ of 3.0 ± 1.0 x 10^-4^ s^-1^ and a *K*_D_ value of 2 nM (23). Stopped-flow kinetics imply that avibactam reacts covalently with TEM-1, a class A β-lactamase (UniProt P62593) within 250 ms (22).

Decarbamoylation/off rates are slow for DBO complexes of SBLs (22, 23); two pathways contribute, either recyclization leading to release of active inhibitor (**Figure 1C**), or hydrolysis, yielding an inactive form (8, 22). Hydrolysis, observed for the carbapenem-hydrolyzing class A SBL KPC-2 (UniProt Q9F663), likely only occurs after on-enzyme desulfation of the bound ligand (22, 23), which as observed by mass spectrometry occurs after 4 hours incubation of DBOs with enzyme (10). The 5-exo-trig recyclization mechanism (**Figure 1C**), favored over hydrolysis in CTX-M-15 (22, 23), is likely promoted when the ring-opened carbamoyl complex is in a stereoelectronically favored conformation, with N6 and C7 in close proximity (**Figure 1C**) (19, 24) and the angle of attack (referred to as the Bürgi-Dunitz angle (α_BD_) (25)) is close to optimal for nucleophilic reactions with a carbonyl group (i.e. ∼105° (26)). The reaction is likely facilitated by the side chains of the conserved catalytic residues Glu166, and particularly Lys73, remaining neutral in the carbamoyl complex (19–21). Recyclization has been evidenced through the use of acyl-transfer mass spectrometry and NMR experiments, which demonstrated avibactam transfer from the CTX-M-15 carbamoyl-enzyme to TEM-1 (and vice versa), at low levels within 2 minutes, and at increasing levels over time (22, 23).

For decades, visualizing enzyme-ligand interactions has largely relied on forming equilibrium complexes with single crystals, trapping these with rapid cryo-cooling, and collecting traditional, rotation-based diffraction data at 100 K. In contrast, time-resolved serial crystallography seeks atomic-resolution structures, at multiple time points, of enzymes engaged in catalysis. Time-resolved serial synchrotron crystallography (tr-SSX) and serial femtosecond crystallography (tr-SFX) at X-ray free electron lasers (XFELs) couple sample delivery methods with reaction initiation strategies that include mixing a microcrystalline enzyme slurry with ligand(s). After a delay time (Δt) for reaction, a still diffraction pattern is collected from each of roughly 10,000 microcrystals (27–29) and all such images merged into a single data set. These diffraction data can produce structural “snapshots” across the reaction coordinate of reactions at temperatures closer to physiological (27). Our previous work with CTX-M-15 microcrystals has shown that an on-demand drop-on-drop approach could resolve, at 1.55 Å resolution (PDB 7BH5), the binding of a β-lactam to CTX-M-15 to form an acyl-enzyme intermediate at Δt = 2 s (29).

Here, we report on use our recently-described drop on fixed target methodology (Kamps *et al*., (2025), BioRxiv), adapted from that developed by Mehrabi and collaborators (30, 31), in which a compact piezo electric dispenser adds 40 - 90 pL ligand droplets onto microcrystals within wells in a silicon chip, which are then probed with X-rays. Importantly, our sample delivery protocol involves ligand addition to alternate wells on the chip in a checkerboard pattern, so providing an internal control for well cross-contamination that is interleaved with the Δt data. We explore the reaction of CTX-M-15 microcrystals with avibactam. We present crystal structures determined from serial data sets collected at both synchrotrons and XFELs using the drop on fixed target system, as well as those obtained from single-crystal experiments on frozen crystals and from room-temperature data collected on microcrystal slurries premixed with ligand, that collectively provide a detailed description of avibactam bound to CTX-M-15. These studies show that avibactam is covalently reacted with CTX-M-15 0.080 s after mixing, and that its conformation at Δt = 1.3 s is distinct from that observed in structures determined from both single-crystal data at 100 K at Δt = 10 s, and from serial room temperature data at Δt = 600 s. Analysis of these structures with molecular mechanics molecular dynamics (MM MD) and quantum mechanics/molecular mechanics (QM/MM) calculations at the density functional theory (DFT) level show that the ligand equilibrates to a state that more rarely samples the early timepoint conformations. These data show that room-temperature serial crystallography data can capture ligand dynamics in an enzyme active site, and is consequently a useful technique for enzyme inhibitor development.

## Methods

### Crystal preparation

Recombinant His-tagged CTX-M-15 (UniProt ID G3G192) was produced, purified and crystallized as described (10, 29). Briefly, CTX-M-15 (expressed in SoluBL21 (DE3) *E. coli*) was purified by mixing with Ni-NTA resin in 50 mM HEPES (pH 7.5) and 400 mM NaCl and eluted with addition of 400 mM imidazole. Following dialysis with 50 mM HEPES (pH 7.5) and 200 mM NaCl, the His-Tag was cleaved with 3C protease (overnight at 4 °C). Following a second passage over Ni-NTA resin, the protein solution was concentrated and loaded onto a Superdex S75 size-exclusion column equilibrated with 50 mM HEPES (pH 7.5) and 150 mM NaCl. Peak fractions were analyzed using SDS-PAGE and pooled before concentrating (10 kDa molecular weight cutoff, Sartorius) to 20 mg mL^-1^.

Macrocrystals were grown using sitting drop vapor diffusion as reported (10). Macrocrystals (50-100 x 100-200 x 200-300 µM) were soaked by addition to a drop containing 12.5 mM avibactam in crystallization solution (2.0 M (NH4)_2_SO_4_, 0.1 M Tris pH 8.0) supplemented with 20% v/v glycerol, before cryo-cooling in liquid nitrogen. CTX-M-15 microcrystal slurries were generated as reported (29). Briefly, the nucleation of CTX-M-15 crystal slurry required the use of seed (generated from crushed macrocrystals). Microcrystal drops comprised 2 μL of crystallization solution (2.0 M (NH4)_2_SO_4_, 0.1 M Tris pH 8.0), 1 μL of seed stock and 2 μL of enzyme (20 mg mL^−1^) and were equilibrated against 500 μL of crystallization solution. Rod-shaped crystals grew within 24 h, with a maximum width of 3 - 8 µm and length of 10 – 20 µm. Crystal density was ∼1 ⋅ 10^8^ crystals mL^−1^. We have shown (29) that these CTX-M-15 microcrystals diffract to high resolution at room temperature, both at Diamond Light Source when mounted on nylon mesh-based fixed target chips (1.65 Å, beamline I24, PDB 7BH6) and at the SACLA XFEL source using a tape drive system (1.6 Å, PDB 7BH3). Pre-soaked serial room temperature datasets were obtained by mixing slurry 1:1 with 50 mM avibactam dissolved in crystallization buffer.

### Drop on fixed target setup and installation

We used a drop on fixed target method for ligand delivery to chip-mounted microcrystals, described in detail in Kamps *et al.,* (2025, BioRxiv). A schematic of the setup is shown in **Figure S21A**. Briefly, CTX-M-15 microcrystal slurries are loaded on to a fixed target silicon chip comprised of 25,600 wells (32), based on chips developed by ourselves and others (33, 34). The chip covers are modified to enable mixing experiments, so that only one side is protected with a 6 µm mylar sheet, while the opposite side is open to allow ligand droplets to be added onto the microcrystals sitting loaded in the wells, similar to previous work (32). 40 µL of microcrystalline slurry suspended in crystallization buffer (∼1 ⋅ 10^8^ crystals/mL, determined using a TC20 cell counter (BioRad), and Neubauer cell counter) is loaded on to the chip in a humidity-controlled environment (>80% relative humidity); vacuum suction is applied to gently remove excess mother liquor.

Ligand is added to every other well using a 50 µm inner diameter piezoelectric injector (PEI), autodrop pipette (AD-KH-501-L6, microdrop Technologies GmbH). To minimize dehydration during data collection, a film cover with damped pads is applied against the open-faced chip. The cover is stationary and includes a 5 × 5 mm^2^ hole that is aligned to the X-ray interaction region, through which ligand droplets are ejected onto microcrystals. The PEI dispenses 40 - 90 pL droplets at ∼1 m/s to every other well, yielding a checkerboard pattern of “+ligand” and “-ligand/control” that consequently interleaves Δt tr-MX and ground-state control datasets across each chip (**Figure S21B**).

### Data collection

For chip-mounted microcrystals, X-ray diffraction data were collected, at room-temperature, from every well of the chip, irrespective of ligand addition, totaling 25,600 images for each chip. Synchrotron data were collected at beamline I24 of Diamond Light Source. Beam size was 7 × 7 µm^2^, with images collected on the Pilatus3 6M detector set at a distance of 320 mm.

PAL-XFEL data were collected on beamline NCI, with an energy of 9.5 keV (pulse energy 650 µJ), and beam size of 3 × 3 µm^2^. Images were collected on the Rayonix MX225-HS detector, set at a distance of 120 mm (∼1.77 Å inscribed circle) and a repetition rate of 30 Hz.

Classical X-ray diffraction data for cryo-cooled macrocrystals were collected at 100 K on beamline I03 of Diamond Light Source, with a Pilatus3 6M detector and an energy of 17.7 keV.

### Data processing and refinement

Serial data were integrated, scaled and merged using Dials (35, 36). For the checkerboard assays, integrated data are split into ‘odd’ and ‘even’ datasets based on image number, representing wells into which ligand was introduced (‘+ ligand’) and wells for which ligand addition was omitted (‘control’), and subsequently scaled and merged separately. This results in a total of 12,800 images for each of the ‘control’ and ‘+ ligand’ datasets across each chip. As required, datasets collected from separate chips were merged. Resolution cutoffs were based on a multiplicity >10 and a monotonic decrease in the CC_1/2_ value. Data quality and completeness were further verified through calculation of omit maps after refinement with the side chain of a bulky aromatic residue (Tyr240) removed, and inspection of *F*_o_-*F*_c_ maps for appearance of electron density consistent with a tyrosine sidechain (**Figure S2**).

X-ray diffraction data from macrocrystals collected at 100 K were integrated, scaled and merged in Dials (35, 36).

Phases were calculated using Fourier synthesis in Phenix with PDB 7BH6 (a room-temperature CTX-M-15 structure determined from serial data collected on a fixed-target chip on DLS I24 (29)) or PDB 6QW8 (a 100 K macrocrystal structure (10)) as the initial models for the room-temperature and 100 K datasets, respectively. In each case the active site SO_4_ ^-^ and ligands were removed. Models were iteratively built in WinCoot (37) with rounds of refinement in phenix.refine (38), with a minimum of 10 rounds to determine ligand and/or SO_4_ ^-^ and solvent occupancy, B-factors and real-space correlation coefficient (RSCC) values. Geometry restraints for non-covalent, intact avibactam (FYG) and covalently bound, ring opened avibactam (NXL) were calculated by eLBOW in Phenix. Omit and isomorphous difference maps were generated in Phenix (38). Figures were generated using PyMOL (39).

### Quantum Mechanics/Molecular Mechanics geometry optimization

Crystal structures from various temperatures and ligand mixing times - 1.3 s and 600 s room temperature structures, 10 s cryo-cooled structures (one for each modelled avibactam conformation) and a 24-hour cryo-cooled structure - were used for Quantum Mechanics/Molecular Mechanics (QM/MM) calculations), following our previously described protocol (40). Protonation states of titratable residues at pH 7.4 were determined using PropKA 3.1 (41), with the exceptions of Glu166 and Lys73, which were simulated in their neutral states as previous computational, NMR and structural studies indicated these residues to be predominantly neutral in covalent avibactam-derived carbamoyl complexes (19–21). Histidine tautomers were determined by the Reduce program of AmberTools23 (42) implemented in AMBER24 (43). The AMBER all-atom ff14SB forcefield was used to describe the protein residues, whilst the generalized AMBER forcefield (GAFF) was employed to describe the carbamoylated avibactam adduct. Each system was solvated in a TIP3P water box, whilst retaining crystallographic waters, with 10 Å separation between solute atoms and the box edge. A 1000 step (300 steps of steepest descent and 700 steps of conjugate gradient) minimization, 25 ps NVT heating (to 298 K using the Langevin thermostat with a 1 ps^-1^ collision frequency) and 500 ps NPT equilibration (using the Berendsen barostat) was undertaken on each structure, with 500 kcal/mol/Å^2^ restraints on all solute heavy atoms applied at each step. This equilibrates the bulk solvent water distribution within the protein crystal, without solute atoms deviating far from their crystallographic positions.

After bulk solvent relaxation was complete, an active site solvent shell was constructed by retaining only the 500 water molecules closest to the carbamoylated avibactam molecule. Each structure underwent a QM/MM geometry optimization, using the Py-ChemShell package (44), employing ORCA 6.0.0 as the QM program (45). Initial optimization was with the carbamoylated avibactam (including the Ser70 side chain) in the QM region, and all atoms within 6 Å of the QM region were included in the active region (atoms that are not frozen and thus can be moved during the geometry optimization calculations – those not in the active region have their positions fixed). The density functional theory (DFT) level of QM description was applied for these geometry optimization calculations, using the B3/LYP functional and 6-31G* basis set with Grimme’s D3 dispersion corrections and Becke-Johnson damping (D3BJ). These optimized structures were used in a second round of geometry optimization where the Ser130, Lys234 and, Lys73 side chains were additionally included in the QM region, with the B3/LYP+D3BJ 6-31G* level of theory. An overview of these fixed and active MM and QM regions is shown in **Figure S3**. Frequency calculations at the geometry optimization level of theory confirmed each structure had minimized to a true minimum.

### Molecular Dynamics simulations

Room temperature crystal structures from the 1.3 s and 600 s mixing timepoints, in which the avibactam *N*-sulfate nitrogen (N6) points away from (N6 ‘out’ conformation’) or towards the C7 (N6 ‘in’ conformation), respectively, were used as starting structures for Molecular Mechanics (MM) Molecular Dynamics (MD) simulations in AMBER24 (42, 43). Crystallographic water molecules were retained and protonation states of titratable residues at pH 7.4 were determined as above, with the exception of Glu166 and Lys73 being neutral (19–21). The N-terminus was protonated. Using tleap (AMBER24), hydrogen atoms were added and systems solvated using a TIP3P (46) octahedral water box with edges 10 Å from protein residues. Overall charges were neutralized with the addition of 3 Cl^-^ atoms. Avibactam-Ser70 carbamoyl complexes were parametrized using the General Amber Force Field (GAFF) parameters in antechamber (47) and restrained electrostatic potential (RESP) fitting to derive partial atomic charges as implemented in the RED Server (48). The ff14SB MM forcefield (49) was used for standard protein residues. First, water oxygens, all hydrogens and ions were minimized (300 steps of steepest descent followed by 700 steps of conjugate gradient) with restraints (500 kcal/mol/Å^2^) on all other atoms, followed by additional minimization of protein sidechains with the same protocol (restraining only backbone atoms). Systems were then heated to 298 K over 50 ps of simulation using a Langevin thermostat (with C_α_ atoms and the avibactam-Ser70 carbamoyl ligand restrained at 100 kcal/mol/Å^2^), then system pressure was equilibrated to 1.0 bar using the Berendsen barostat over 3 × 250 ps, with restraints on the C_α_ atoms and the avibactam-Ser70 carbamoyl ligand (restraint weights were decreased from 50 kcal/mol/Å^2^ to 5 kcal/mol/Å^2^). A final 250 ps equilibration step was run with 2 kcal/mol/Å^2^ restraints on the avibactam-Ser70 carbamoyl ligand only, to moderately constrain the ‘in’ and ‘out’ conformations of the *N*-sulfate nitrogen in the 600 s and 1.3 s structures (**Figure S20**), respectively, prior to production runs. Structures were then simulated using MM MD for 500 ns in triplicate (a total of 1.5 µs for each structure). Simulations were analyzed in CPPTRAJ (50) in Amber 24. Root mean square deviation (RMSD) calculations used the final equilibrated structure prior to the production runs as the reference.

## Results

### Room temperature drop on fixed target mixing resolves avibactam covalently bound to CTX-M-15 at 1.3 seconds

To obtain room-temperature time-resolved serial crystallography data, we used our drop on fixed target setup (**Figure S21**), in which the reaction is initiated by adding picolitre ligand droplets onto crystals within wells (Kamps *et al*., (2025), BioRxiv). Synchrotron [DLS I24 (UK)] and XFEL [PAL-XFEL (South Korea)] diffraction data were collected for uncomplexed/resting crystals and from crystals after 1.3 s exposure to 200/300 mM avibactam (265 Da), which is highly water soluble, delivered through ejection from a piezoelectric injector (**Table S1**). Data extended to 1.62 Å - 1.92 Å resolution at DLS, and 1.45 Å - 1.60 Å at PAL-XFEL. Comparison of ‘resting state’ room temperature structures with a structure determined from a diffraction dataset from a cryo-cooled (100 K) single crystal grown under the same conditions [PDB 4HBT (51), 1.1 Å resolution] (**Figure S4**) shows moderate differences in active site architecture. In particular, there is movement of an active site water molecule responsible for deacylating β-lactam complexes (DW, **Figure S4**) closer to the general base (Glu166) in the room temperature structures. Further, Lys73 forms a Lys73 - Ser130 hydrogen bond in the structures determined at room temperature, or a Lys73 - Asn132 hydrogen bond in the structure determined at 100 K.

For the drop on fixed target experiments, we implemented important internal controls to assess the possibility of ligand contamination across wells in the chip (described in more detail in Kamps *et al*., (2025), BioRxiv). To this end, 200/300 mM avibactam was ejected into alternate wells of a chip previously loaded with CTX-M-15 microcrystals, and Δt = 1.3 s diffraction data split into ‘+ligand’ and ‘control’ (i.e. images from wells to which avibactam had not been added) datasets. Visual inspection of the resulting electron density maps, including *F*_o_ - *F_c_*from ‘+ligand’ and ‘control’ datasets, isomorphous *F*_o(+ligand)_ - *F*_o(control)_, and isomorphous *F*_o(+ligand)_ - *F*_o(resting)_ (**Figure 2A** and **Figure S5**), revealed that avibactam had reacted to form the covalent carbamoyl ester to Ser70 (NXL in **Figure 2A**), and further showed that ligand ejection had not resulted in contamination of adjacent wells (*F*_o(control)_ - *F*_o(resting)_, **Figure S5**).

**Figure 2.**
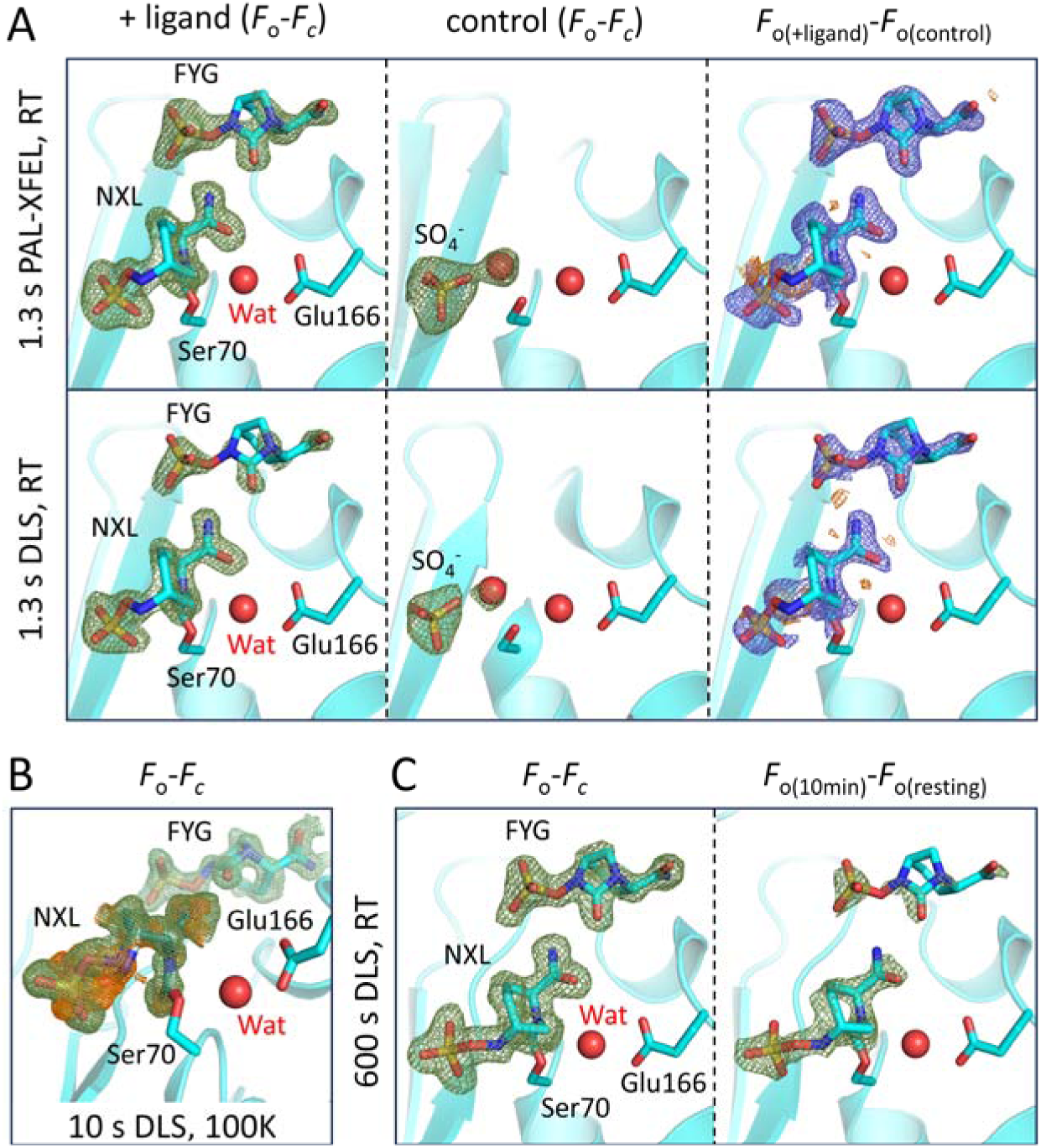
Avibactam reaction with CTX-M-15 over time from room temperature and cryo crystallography. Views from the active sites of avibactam bound in CTX-M-15 (NXL is the covalently bound avibactam, and FYG is the non-covalent, intact avibactam bound at the entrance to the active site). Electron densities (mesh) are contoured at 3σ around avibactam molecules or the active site sulphate and water. **(A)** Room temperature serial crystallography after 1.3 s soaking using the fixed target drop on chip approach at PAL-XFEL (top) and I24 (bottom). *Left*, *F*_o_-*F*_c_ electron density (green), from data collected from wells in which ligand was ejected (+ ligand). *Middle*, *F*_o_-*F*_c_ electron density (green), in control wells in which ligand is not ejected (control). *Right*, isomorphous difference maps (*F*_o_-*F*_o_) between +ligand and control, contoured at +(blue)/-(orange) 3σ. **(B)** 10 s soak using macro crystals at 100 K. 2 conformations of the covalently reacted avibactam are observed. *F*_o_-*F*_c_ electron density is shown after removal of each conformation (coloured green for conf A and orange for conf B). **(C)** Room temperature serial crystallography structure of CTX-M-15 microcrystals after 10-minute exposure to avibactam and loading on to fixed target chips. *Left*, *F*_o_-*F*_c_ density (green); *right*, isomorphous *F*_o(10min)_-*F*_o(resting)_ (green).

In structures determined from data collected at PAL-XFEL and DLS, the covalently bound avibactam ligand (NXL) was refined at full occupancy, with RSCC values of 0.96 and 1.00, respectively (**Table S2**). A second DBO intact, bicyclic avibactam molecule (FYG, **Figure 2**) is also observed non-covalently bound at the entrance to the active site, with occupancies/RSCCs of 0.66/0.91 and 0.62/0.99 for the PAL-XFEL and DLS data, respectively (**Table S2**). A PAL-XFEL structure after 0.080 s avibactam exposure was also consistent with formation of a covalent complex between avibactam and Ser70 (Kamps *et al*., (2025) BioRxiv); however, ligand electron density was less well defined compared to the Δt = 1.3 s structures (**Figure S6**). Therefore, the bound avibactam conformation could not be definitively determined, likely due to incomplete ligand occupancy (33%). Although shorter timepoints are possible using this initiation method, early intermediates in the CTX-M-15+avibactam+Δt reaction (e.g. Michaelis complex, tetrahedral intermediate, or initial ring-opened state) are very likely short-lived and were not observed. Key limitations preventing observation of such states likely include the relatively long time required for avibactam to diffuse through the crystal lattices and any buffer remaining in the fixed target chip wells, coupled with the faster kinetic steps that precede the stable, covalent complex.

Tr-SSX data (Δt = 1.3 s) collected at DLS, with avibactam at concentrations of 20, 50 and 300 mM in the piezoelectric injector (**Table S3**), indicated that ligand concentration impacts on the refined occupancy of the covalently bound avibactam (NXL) and of the intact molecule at the active site opening (FYG) (**Figure S7**). Full NXL occupancy, and partial FYG occupancy (62%) was observed in the structure determined after addition of 300 mM avibactam. With 200 mM avibactam (PAL-XFEL Δt = 1.3 s dataset), there was little difference in the final refined RSCC or adjusted B-factor for NXL compared with those for structures obtained with 300 mM avibactam in the ejector (**Table S2**). Addition of 50 mM avibactam yielded a structure with NXL at 78% refined occupancy, with an RSCC of 0.94, but with no modellable density for FYG, with isomorphous *F*_o(+ligand)_-*F*_o(control)_ maps showing disjointed electron density in this region. Using 20 mM avibactam, we did not obtain evidence for either NXL, or the presence in the active site of intact (i.e. unreacted) avibactam, with only very minor differences in the *F*_o(+ligand)_-*F*_o(control)_ isomorphous electron density maps (**Figure S7**). These results indicate that the milliseconds to seconds equilibration times in our tr-SSX and tr-SFX methods require higher ligand concentrations compared to traditional cryo-crystallography with single macro-size crystals and prolonged equilibration times.

We next investigated the relative stability of the second, DBO intact avibactam molecule (FYG) observed at the active site opening, with a series of ‘wash out’ experiments using macro-size crystals and traditional data collection methods. CTX-M-15 crystals were soaked for 3 hours with 12.5 mM avibactam, then manually transferred (in loops containing ∼5 nL well reagent) into wells containing 10 μL crystallisation reagent supplemented with 20% glycerol, but without avibactam, before cryo-cooling in liquid nitrogen at prescribed times. The resulting diffraction data extended to 0.87 – 1.22 Å resolution (**Table S4**). After 10 s and 50 s in the ‘wash out’ buffer, FYG could still be observed at high occupancy and RSCC, while at longer time points (600 – 1800 s) was washed out of the crystals, shown by comparative isomorphous maps calculated against uncomplexed CTX-M-15 (PDB 4HBU (51)) (**Figure S8**). However, the covalently bound NXL was retained throughout. Therefore, the second, intact, avibactam equivalent (FYG) likely represents an artifact of the high avibactam concentrations required for these rapid mixing experiments, that accumulate *in crystallo*.

### Time-resolved movement of the carbamoylated avibactam

In addition to the drop on fixed target data, which yielded structures after exposure to avibactam over relatively short (ms – s) timescales, we explored longer time points through pre-soaking experiments. These used both single-crystal cryo crystallography and tr-SSX; data were collected at 100 K from macrocrystals looped and cryo-cooled after ∼10 s exposure to 12.5 mM avibactam (**Table S4**), as well as at room temperature from microcrystals equilibrated with 25 mM avibactam at Δt = 300 – 600 s (300 s presoaking, followed by 300 s data collection, from the same chips used for the drop on fixed target experiments) (**Table S5**). As with the Δt = 1.3 s data, *F*_o_-*F*_c_ omit electron density maps (calculated after ligand removal) revealed the presence of both a molecule of covalently bound ring-opened avibactam (NXL) and a molecule of non-reacted intact avibactam (FYG) in the CTX-M-15 active site in both the Δt = 10 s cryo and Δt = 600 s room temperature structures (**Figure 2B** and **Figure 2C**, respectively). Of note, a previous structure of a CTX-M-15:avibactam complex, determined from diffraction data collected after >24 h co-crystallisation with 10 mM avibactam [PDB 4HBU (51)], does not contain FYG. In the Δt = 10 s cryo structure reported here, 2 NXL conformations (A and B) were observed, with refined occupancies of 0.33 and 0.67 (**Table S2**), differing in the positioning of the *N*-sulfate moiety (**Figure S9)**.Of note, room-temperature data shows differences in active site hydration, with the position of the active site water molecule proposed to be involved in the decarbamoylation reaction consistently 0.2 – 0.3 Å further away from the avibactam C7 in the room temperature compared to the cryo structures (**Figure 3**).

**Figure 3.**
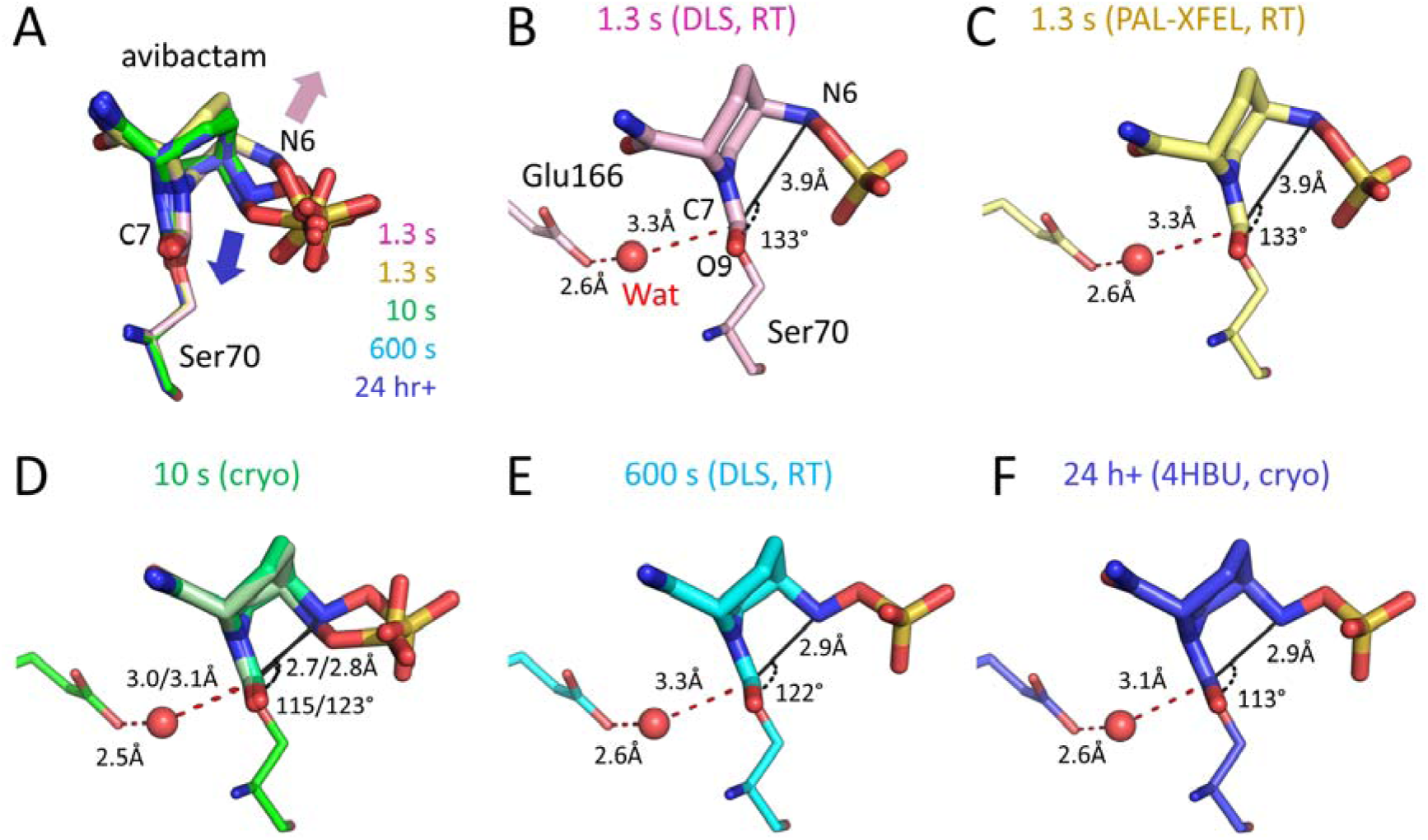
Time-resolved movement of the avibactam N6 nitrogen. **(A) Active site views of avibactam reacted with CTX-M-15 Ser70.** 1.3 s (DLS, pink C atoms; PAL-XFEL, yellow), 10 sec (green C atoms), 600 s (cyan C atoms) and co-crystal (blue C atoms, PDB:4HBU (51)) structure views are superposed. Arrows denote variation in nitrogen position across these datasets, either pointing ‘in’ towards the C7 carbonyl (blue arrow), or ‘out’, away from the C7 carbonyl (pink arrow). **(B-F) Key measurements of the time-resolved carbamoyl complexes.** Shown are the N6 to C7 ‘recyclization’ reaction distance and N6-C7-O9 angle, and distances between the presumptive decarbamoylating water (Wat) and Glu166 or the C7 of reacted avibactam. **(B) 1.3 s room temperature Diamond Light Source (DLS)** dataset. **(C) 1.3 s room temperature PAL-XFEL** dataset. **(D) Cryo (100 K)** dataset at DLS from which the refined model converged with two partial occupancy ligand conformations, **(E) 600 s room temperature** dataset collected from fixed target chips at I24 (DLS), **(F) 100 K co-crystal** (PDB 4HBU).

In structures determined at all Δt timepoints, NXL interacts with approximately eight residues around the active site (**Table S6 and Figure S10**). In particular, there are hydrogen bonding interactions with the side chain of Ser130 (via the exocyclic N6), the side chain nitrogen atoms of Asn104 and Asn132 (via the amide oxygen O11) and the side chain hydroxyl groups of Ser130, Thr235 and Ser237 (with the O14/15/16 oxygens of the sulfate moiety). The C7 carbonyl oxygen (O9) sits in the oxyanion hole formed by the backbone amides of Ser70 and Ser237. FYG is situated at the opening of the CTX-M-15 active site and has not been previously observed in SBL:avibactam complexes. The FYG sulfate forms weak interactions with NXL-NH_2_, and its C7 carbonyl is within hydrogen bonding distance of the side chain nitrogen of Asn104 (**Table S6**). However, a comparison of the 10 s and 1800 s ‘wash’ structures (described above) indicates that loss of FYG has no impact on the binding or positioning of the covalently reacted avibactam (NXL, **Figure S11** and **Table S6**).

The most significant difference observed across the different datasets relates to the positioning of the *N*-sulfate nitrogen (N6) of NXL, which adopts an ‘in’ or ‘out’ conformation (blue and pink arrows in **Figure 3A**) with respect to the C7 carbonyl carbon of the carbamoyl ester link to Ser70. At Δt = 1.3 s, in both DLS and PAL-XFEL data, N6 points ‘out’, away from the C7 carbon, with N6 - C7 distances of 3.9 Å. The N6-C7-O9 angles (the α*_BD_*) are 133°, not close to the optimal of ∼105° for recyclization (**Figure 3B** and **3C**). The high resolution Δt = 10 s cryo dataset refined to convergence with an atomic model for the covalently-attached NXL in two, partially occupied conformations (0.33/0.67, **Table S2**); both show an inward shift of N6 (**Figure 3D**), but with the sulfate present in two conformations. Compared to the 1.3 s structures, both configurations have shorter N6 - C7 distances (2.8 (A)/2.7 Å (B)) and more acute N6-C7-O9 angles (115° (B), 123° (A)), suggesting flexibility and instability at this region at Δt = 10 s. The Δt = 600 s dataset yields a structure resolving only the ‘in’ conformation (**Figure 3E**), and is most similar to the previously published >24 h cryo structure (PDB 4HBU, **Figure 3F** and **Figure S12**). The N6 - C7 distance in both these structures stabilizes at 2.9 Å, although the N6-C7-O9 angle decreased from 122° to 113° between the 600 s and 24 h structures, respectively. These movements, observed over time points ranging from 1.3 s to 24 h, are further reflected in the distance from N6 to the Oγ of residue Ser130 (**Table S6 and Figure S13**); Ser130^Oγ^ is proposed to be involved in abstracting the proton from the *N*-sulfate nitrogen (N6) to enable DBO recyclization (19) (**Figure 1C**). In the Δt = 1.3 s structures, with N6 in the ‘out’ conformation, the N6 - Ser70^Oγ^ distance is 3.3/3.4 Å (**Figure S13B** and **Figure S13C**). This shortens over time because of movement of N6, with the flexibility observed in the structure at Δt = 10 s resulting in distances of 2.4 (conf. A) and 2.9 Å (conf. B) (**Figure S13D**), which then equilibrate in the 600 s and 24 h ‘in’ structures to 3.0 and 2.9 Å, respectively (**Figure S13E** and **Figure S13F**). Conformation B in the Δt = 10 s structure therefore most closely represents the Δt = 24 h ‘in’ conformation, while conformation A appears to be a relatively flexible intermediate state that relates to neither ‘in’ nor ‘out’ conformations.

### Computational modeling of avibactam dynamics and conformations

The positions of hydrogen atoms are particularly important for recyclization of the carbamoylated avibactam, with proton transfers involved in the reaction mechanism (19), but these cannot be defined in our electron density maps. We employed QM/MM geometry optimization to position hydrogen atoms, and thus identify species corresponding to energy minima across the singly protonated N6 ‘out-in’ transition of the carbamoylated complex. We analyzed five different atomic models derived from the datasets collected at different Δt timepoints: 1.3 s room-temperature; 10 s room temperature equilibration with 100 K data collection (conformations A and B); 600 s room temperature; 24 h room temperature equilibration with 100 K data collection [PDB ID 4HBT (51)]. In all five optimized structures (**Figure 4**), key residues involved in hydrolysis and recyclization (Ser130, Ser70, Lys73, Glu166) retained the same positions (**Figure 4A**); in particular, the Ser130 γ-hydrogen points to the lone pair on Lys73 (1.7 Å distance). Furthermore, the water molecule (Wat, **Figure 4**) potentially positioned for hydrolysis of the avibactam carbamoyl, is in approximately the same location, relative to the C7 carbon and Glu166^Oε1^. The 1.3 s room temperature optimized structure maintains the N6 ‘out’ conformation (i.e. it does not move to the ‘in’ conformation), indicating this is at a local energy minimum, with the avibactam N6 hydrogen directed 4.4 Å away from Ser130^Oγ^ (**Figure 4B** and **Table S7**) but with N6 lone pair and proton interacting with the aromatic ring of Tyr105. The optimized structures starting from the 10 s (conformation B), 600 s and 24-hour ligand soaking experiments all represent an ‘in’ conformation, in which N6 and its hydrogen have rotated inwards to form a hydrogen bond with the Ser130^Oγ^ (distance 1.9 Å, **Table S7**), and no longer interact with Tyr105. There were minor differences between these three optimized structures in the N6-C7-O9 angle (α*_BD_*), H-C5-N6-H dihedral and C7 - N6 distance (108° - 110°, 301° - 307°, 2.5 - 2.9 Å, respectively, **Table S7**), but all maintained equivalent ‘in’ conformations primed for recyclization with α*_bd_*close to its 105° optimum. In the QM/MM optimized structure derived from conformation A of the Δt = 10 s structure, the N6-hydrogen has rotated to hydrogen bond with Ser130 (1.9 Å) and no longer interacts with Tyr105. However, the N6-C7-O9 angle (α*_BD_*) of the carbamoylated avibactam (117.2°), the H-C5-N6-H dihedral (280.5°), and the C7-N6 distance (3.5 Å) all indicate that the N6 nitrogen is at a point midway between its position in the structure at Δt = 1.3 s, and its position in all other structures, but is not yet primed for recyclization.

**Figure 4.**
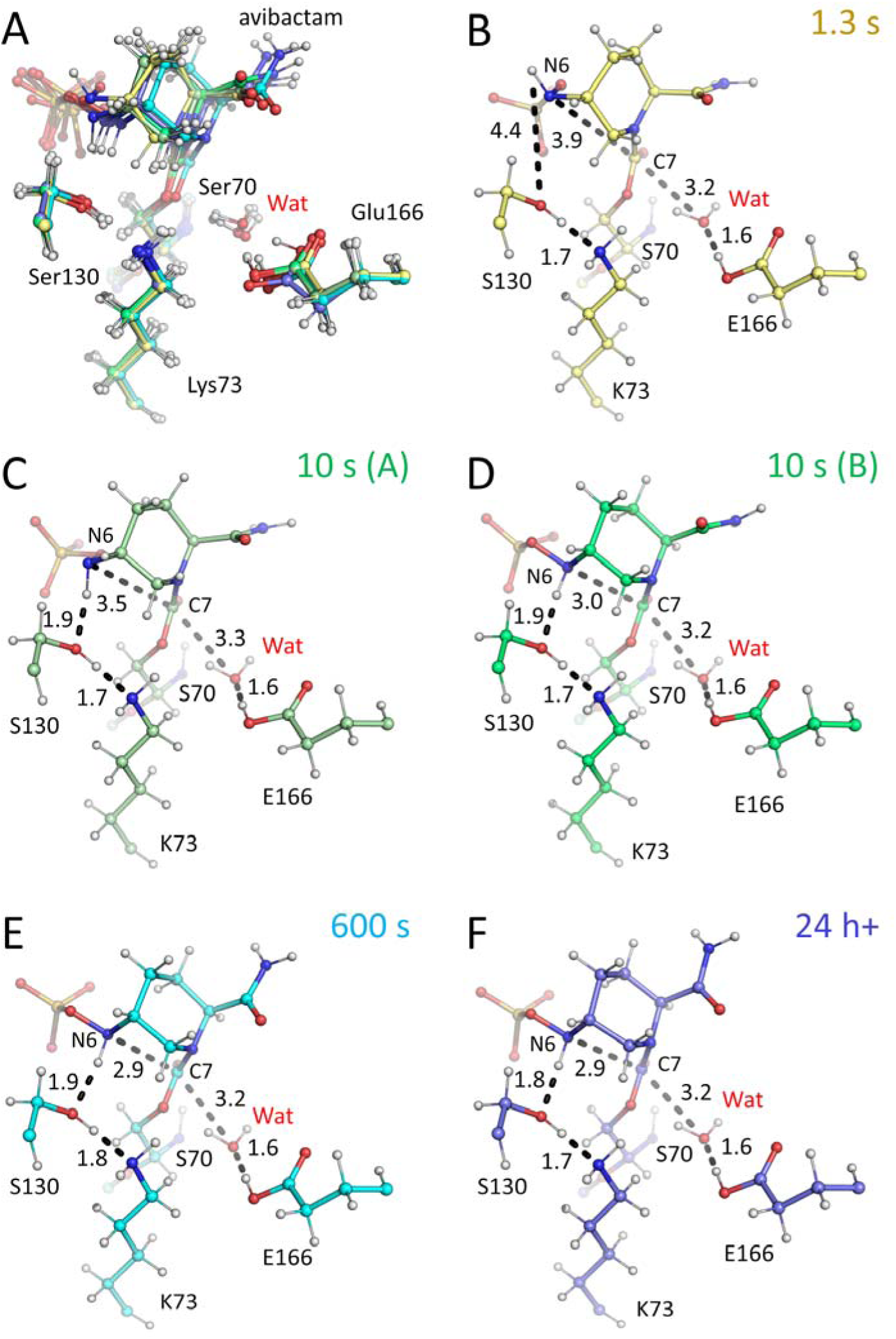
QM/MM optimised active site models of carbamoylated complexes of CTX-M-15. QM/MM at the DFT level was used to optimise CTX-M-15 X-ray structures (overlay in **A)** after avibactam exposure for **(B)** 1.3 s room temperature PAL-XFEL data (N6 ‘out’ conformation, yellow), **(C)** 600 s cryo data, conformation A (N6 ‘out’, pale green), **(D)** 600 s cryo data, conformation B (N6 ‘in’ conformation, bright green), **(E)** 600 s room temperature (N6 ‘in’, cyan), **(F)** 24 h+ cryo (N6 ‘in’, blue). The side chains of key catalytic residues are shown, and distances (black dashes) are labelled in Å. Note, neutral protonation states of Lys73 and Gly166 are based on prior work (19–21), as described in the main text.

To further investigate the implications for the dynamics of the respective avibactam complexes of our structures determined by time-resolved X-ray crystallography, that represent discrete ‘snapshots’ along the reorganization pathway, we used MM MD to simulate the N6 ‘out’ and ‘in’ conformations directly from the room-temperature structures determined from the Δt = 1.3 s high-resolution PAL-XFEL, and Δt = 600 s DLS data sets, respectively. In each case, three replicate 500 ns simulations were carried out. The second, non-covalently bound, avibactam molecule (FYG) in the structures was omitted from the calculations, as this is most likely an artifact of the crystallography at high avibactam concentrations (see above). The simulations remained stable and equilibrated over the length of the trajectories and did not show major differences in the movements of backbone atoms (as defined by root mean square deviation (RMSD) and root mean square fluctuation (RMSF) values, **Figure S14**). Analysis of the dynamics of the carbamoylated avibactam molecule, however, shows significant differences between the simulations starting from the N6 ‘in’ and ‘out’ conformations. In particular, the RMSD values indicated that the covalently-linked avibactam (NXL) in simulations based on the Δt = 1.3 s ‘out’ conformation was more dynamically flexible than in those based on the Δt = 600 s ‘in’ conformation (**Figure 5A**), with average RMSDs of 1.15 Å and 0.92 Å, respectively. Inspection of by-atom RMSF plots shows that this is largely due to mobility of the SO_4_ ^-^, N6 and carbamide, while all other NXL atoms remain relatively stable (**Figure 5B**). To better understand these movements around the *N*-sulfate N6, we plotted the N6 - C7 distances and H-N6-C5-H dihedral angles (**Figure 5** and **Figure S15**). In the simulation based on the Δt = 1.3 s structure the N6 - C7 distance is on average longer than is the case in the simulation based on the Δt = 600 s structure (3.64 Å vs 3.20 Å), consistent with the crystal structures. However, these analyses show that in both simulations avibactam can adopt distinct conformations. In the Δt = 600 s (‘in’) trajectories, 2 major conformations are observed, particularly with respect to the dihedral angle (**Figure S15**), both corresponding to the N6 nitrogen pointing towards the C7, but with differences in positioning of the associated N6. However, a minor cluster indicates this simulation also sampled the Δt = 1.3 s (‘out’) conformation (snapshot 3 in **Figure 5C**), albeit relatively rarely (<1% of the total 1.5 µs of simulation). When starting from the ‘out’ conformation, 3 major conformations were observed, consistent with RMSD and RMSF data that indicated the carbamoyl-avibactam was more dynamic in simulations starting from this conformation than from the ‘in’ conformation. Notably, these Δt = 1.3 s simulations frequently sampled both ‘in’ (see snapshots 1 and 2 in **Figure 5D**) and ‘out’ conformations (snapshots 3 and 4 in **Figure 5D**). The low occupancy flexible conformation A in the Δt = 10 s crystal structure was not sampled in any of the trajectories. Differences in the N6-C7-O9 angle are less pronounced between ‘in’ and ‘out’ conformations, due to movements of the carbamoyl ester linkage, but show a slight increase in this angle in the simulations based on the 1.3 s (‘out’) than the 600 s (‘in) structure, consistent with the crystal structures (**Figure S16**). In addition, the mean Ser130^Oγ^ - Avi^N6^ and Ser130^Oγ^ - Lys73^Nζ^ distances, considered important in the recyclization reaction, trend longer over the course of the Δt = 1.3 s simulations than the Δt = 600 s simulations (3.64/3.20 Å and 3.62/3.34 Å, respectively; **Figure S17A** and **S17B**).

**Figure 5.**
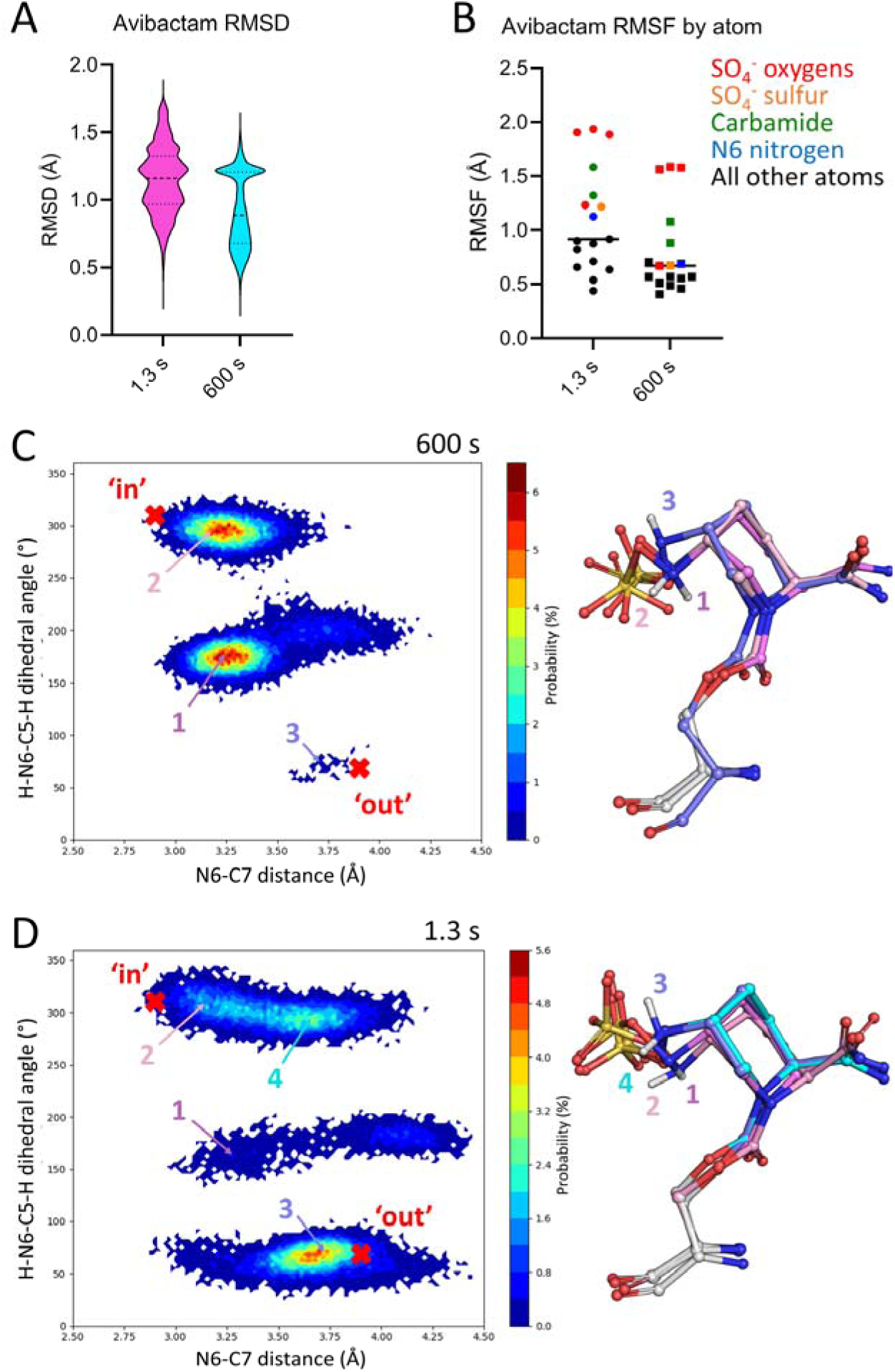
Molecular dynamics analysis of 1.3 s and 600 s room temperature structures. **(A)** Average root mean square deviation (RMSD) of avibactam atoms over the total 1.5 μs simulation time for 1.3 s (pink) and 600 s (cyan) room temperature structures. **(B)** Average root mean square fluctuation (RMSF) of each avibactam atom during the simulations. **(C)** and **(D)** Probability heatmap plots of the distance between avibactam C7 and N6 (in Å) over the course of the 600 s and 1.3 s simulations, respectively, against the H-N6-C5-H dihedral (°). Red crosses indicate values for the 1.3 s ‘out’ and 600 s ‘in’ crystal structures. *Right*, structures from representative frames of the simulations, numbered according to their position on the plot, *left*.

## Discussion

The interaction of clinically relevant drugs (both inhibitors and substrates) with β-lactamases over short time scales is poorly understood, largely due to a lack of such time-resolved structural information. We employed time-resolved serial X-ray crystallography to explore the early stages of binding of the potent small molecule inhibitor avibactam to the β-lactamase CTX-M-15. Contrary to early thinking, in which it was thought that there is little conformational movement of the avibactam molecule on ring opening (23), our work shows that the conformation of avibactam changes over time when covalently bound to CTX-M-15, particularly with respect to the positioning of N6 in relation to the C7 carbamoyl of the carbamyl ester. Two major conformations are observed: an N6 ‘out’ conformation typified by the room temperature Δt = 1.3 s structures, and an N6 ‘in’ conformation captured in the room temperature Δt = 600 s structure. The latter is likely the conformation relevant for recylization (51). These apparently transition *via* a flexible intermediate, potentially corresponding to conformation A in the Δt = 10 s structure and/or its DFT optimized conformation. MD simulations presented here indicate that the N6 ‘in’ and ‘out’ states can interconvert, particularly when starting from the ‘out’ conformation, in which alternative states are frequently sampled. When starting from the ‘in’ conformation interconversion, while a rare event, is still observed. In contrast a previous single 100 ns MD trajectory (20) starting from the ‘in’ conformation only (based on PDB 4HBU (51)), did not undergo movement to the ‘out’ conformation. This suggests that our extensive (1.5 µs) simulations more completely sample the flexibility of the two conformations.

The avibactam carbamoyl complexes of serine β-lactamases can resolve either through recyclization to regenerate intact and active inhibitor or by hydrolysis. In CTX-M-15, decarbamoylation likely proceeds through recyclization rather than hydrolysis, as supported by previous QM simulations that show the calculated barriers for recyclization to be lower than those for hydrolysis (19). Positioning of the N6 nitrogen is required for stereoelectronically favored avibactam recyclization and reformation to give the DBO, as the N6 to C7 distance and α*_BD_* angle are linked to the likelihood of attack by N6 on C7 (**Figure 1C**). Compared to the 1.3 s ‘out’ conformations, in which the bound avibactam atoms are poorly aligned for recyclization, from 10 s onwards, bound avibactam (in the ‘in’ conformation) appears better primed for recyclization, with N6 - C7 distances reducing and α*_BD_* becoming more acute and closer to optimal (**Figure 3**).

NMR and high-resolution crystal structures of CTX-M-14 (80% sequence identity to CTX-M-15, with active site residues conserved) in complex with avibactam indicated that Lys73 and Glu166 are neutral in the carbamoyl complex (21). For recyclization, therefore, proton abstraction from Ser130 by the neutral Lys73 is followed by proton transfer from avibactam N6 to Ser130 (**Figure 1C**). Our QM/MM calculations show that in all our structures, including both ‘in’ and ‘out’ conformations, Ser130 and Lys73 are primed for this first step by forming a conjugate ion pair. However, the avibactam N6 hydrogen is only close enough to Ser130 to support proton transfer in the ‘in’ conformations. While the N6 – Ser130^Ογ^ distance is similarly short in conformation A of the Δt = 10 s structure, the DFT-optimized N6 - C7 distance (3.5 Å) and α*_BD_*angle (117.2 °) indicate it is likely not primed for recyclization.

For avibactam hydrolysis, which presents the alternative route to regeneration of active enzyme, it is likely Lys73 must first abstract a proton from the general base Glu166, that then activates a water molecule (Wat in **Figures 2, 3** and **4**) for nucleophilic attack on the avibactam C7 (19, 21). In our MD trajectories, the distance between these two residues is greater than 3.5 Å in 71% and 91% of the frames for the Δt = 1.3 s (‘in’) and Δt = 600 s (‘out’) structures, respectively (**Figure S17C**), suggesting these are not consistently primed to initiate the hydrolysis reaction. On the other hand, in our experimental time-resolved structures, the α*_BD_* angle of the water to the carbonyl bond increases from 93° in our Δt = 1.3 s structure, to 100° at Δt = 600 s, so closer to the Bürgi-Dunitz optimal for nucleophilic attack, despite little change in the distances to Glu166 and the avibactam C7. However, we note that, in the structure of an avibactam carbamoyl complex of the KPC-2 carbapenemase, for which hydrolysis has been determined to be a more likely decarbamoylation pathway than in CTX-M-15 (23), the general base Glu166 - carbamoylating water (Wat) and Wat – carbamoyl carbon (C7) distance are shorter than their equivalents in CTX-M-15 (CTX-M-15 1.3/600 s: Glu166 – *2.6 Å* – Wat – *3.3 Å* – C7; KPC-2 (PDB 4ZBE (52)): Glu166 – *2.4 Å* – Wat – *2.8 Å* – C7). Therefore, based on data collected at 100 K, and consistent with results from kinetic assays, the KPC-2 system appears better primed for hydrolysis than does CTX-M-15. However, room-temperature KPC-2:avibactam structures will be required to provide a more accurate picture of active site hydration under conditions closer to physiological.

Comparison of our structures with previously determined structures of avibactam carbamoyl complexes of β-lactamases (from single-crystal experiments at 100 K, after exposure to avibactam at varying timepoints), shows some similarities (**Figure S18**). As expected, based on our computational and room-temperature crystallographic data, that show it to be the most stable conformation, the ‘in’ conformation is seen in structures of avibactam complexes of CTX-M family enzymes obtained either through co-crystallisation or from long soaks, i.e. ≥ 24 hours incubation of enzyme with avibactam [CTX-M-15, PDB 4HBU/4S2I (24, 51); CTX-M-151, 6BPF (53), **Figure S18A**)]. To date, the ‘in’ conformation has only been seen in one structure of an avibactam complex of a non-CTX-M family β-lactamase, the carbapenemase CRH-1 [PDB 8EK9 (54), **Figure S18A**]. Although our MD identifies that the ‘out’ conformation can sample multiple conformations in CTX-M-15, it has been observed in multiple other structures of avibactam complexes of class A SBLs determined by single-crystal diffraction of crystals frozen after soaking times ranging from 8 min – 24 hours: e.g. BlaC, 4DF6 (55); CTX-M-14, 6MZ1 (21); KPC-2, 4ZBE (52); KPC-44, 8TMR (56); L2, 5NE3 (57); PenA, 7DOO (58); PER-2, 6D3G (59); SHV-1, 4ZAM (52); TEM-1, PDB 8DE0 (60); TLA-3, 5GWA (61); VCC-1, 6MKQ (62) (selected examples in **Figure S18B**; a sequence alignment of example enzymes from **Figure S18** is shown in **Figure S19**).

Both ‘in’ and ‘out’ avibactam conformations have previously only been observed for CTX-M-14. The ‘out’ state was seen after 1 s exposure to avibactam at pH 4.5, obtained using spitrobot mixing and data collection at 100 K (63). From data collected at 100 K, 1800 s after mixing with avibactam at pH 7.9 (21), three covalently bound avibactam conformations were modelled, with one aligning with the ‘in’ conformation B in the Δt = 10 s, 600 s and 24 h structures (**Figure S18C**); in a Δt = 1800 s cryo complex at pH 5.3 (in the same study), only the ‘out’ conformation was modelled. Finally, in structures of avibactam:CTX-M-14 complexes at pH 6.0 (from data collected at room temperature after a non-specified soaking duration), and pH 7.9 (from co-crystal data collected at 100 K) only the ‘in’ conformation was modelled (PDBs 6GTH (64) and 9OR3 (65)). Although these data suggest there may also be equilibration to the ‘in’ conformation of avibactam over time in the related CTX-M-14, data were collected through different techniques and at different pH values (likely biasing or stabilizing the conformations by changing the protonation state of Lys73) complicating direct comparison.

The observed ‘out’ conformations in the complex of CTX-M-15 carbamoylated with avibactam at 1.3 s must proceed via an ‘in’ type species after ring opening, at which point the avibactam N6 is protonated (likely by Ser130 (66)) before it then ‘flips’ away from the C7 carbonyl carbon (to the ‘out’ conformation). Given we do not observe this intermediate at 1.3 s, it is probably less stable than either the ‘out’ or ‘in’ conformations described by our crystallography. Following formation of the ‘out’ conformation, our computational and crystallographic data indicate this can equilibrate to the ‘in’ over time (both with a singly protonated N6), which is then primed for recyclization. The ‘in’ conformation (with a protonated N6) observed from 10 s onwards is relatively stable in the crystal over time, and so unlikely represents the intermediate species formed immediately after ring opening but prior to outwards movement of N6. Consistent with this, the QM optimized models with a protonated N6 do not substantially deviate from the crystal structures, with the proton remaining bonded to N6 throughout. The mechanism and conformational movements during initial ring opening and N6 protonation that result in the ‘out’ conformation will require more detailed investigation, including extension of QM/MM simulations to study the reaction and proton movements considering these time-resolved data. However, our results are further consistent with, and add further evidence for, recyclization being the preferred route for reversing avibactam inhibition of CTX-M-15.

## Conclusions

Time-resolved methods to investigate enzyme complex structures which require ligand diffusion into their active site are still in early development stages, but full understanding of such reactions requires high-resolution structural descriptions at all stages of the reaction pathway. Serial crystallography strategies are well suited to studying ligand binding, dynamics, and catalysis within microcrystals at near physiological temperatures. Here, we use a drop on fixed target method that has been adapted for use at synchrotrons as well as XFEL sources, enabling collection of high-resolution diffraction data in serial crystallography experiments. This platform has supported the study, at room-temperature, of formation of the covalent complex of the diazabicyclooctane inhibitor avibactam with the CTX-M-15 β-lactamase over time scales from 0.08 s onwards. Our work is facilitated by the rapid diffusion of ligand-containing droplets into enzyme microcrystals, and by our implementation of a data collection protocol that negates the possibility of including images from microcrystals accidentally exposed to ligand during injection into adjacent wells of the target chip. These data reveal the dynamics of a ligand covalently bound to an enzyme active site, and highlight the importance of collecting time-resolved diffraction data at room temperature to obtain more complete understanding of the mechanistic basis of enzyme interactions and reactions with ligands. Uncovering such information, at temperatures closer to physiological, will identify interactions important to formation and maintenance of stable enzyme:inhibitor complexes, so facilitating their optimization in inhibitor (re)design. Such approaches will aid in structure-guided drug discovery processes aimed at identifying new compounds that target β-lactamases and other clinically important nucleophilic enzymes.

## Acknowledgments

We thank Diamond Light Source for beamtime (mx23269, mx25260, mx32727 and mx39247) and the staff of beamlines I24 and I03 for their assistance. The experiments were performed using an NCI instrument at PAL-XFEL (proposal number 2023-2nd-NCI-014, June 2024). The authors would like to thank all the members of PAL-XFEL for their assistance with the experiments and data collection. The authors thank the Global Science experimental Data Hub Center (GSDC) at the Korea Institute of Science and Technology Information (KISTI) for providing computing resources and technical support. All simulations were conducted using the facilities of the Advanced Computing Research Centre at the University of Bristol (http://www.bris.ac.uk/acrc/). This work is part of a project that has received funding from the European Research Council under the European Horizon 2020 research and innovation program (PREDACTED Advanced Grant Agreement no. 101021207) to AJM and JS. AMO, RLO, PA, and JJAGK received support from Diamond Light Source and UK Science and Technology Facilities Council (STFC). These results were supported in part by a Royal Society Wolfson Fellowship RSWF\R2\182017 and a Wellcome Investigator Award 210734/Z/18/Z to AMO. Research was supported by the BBSRC-funded South West Biosciences and Interdisciplinary Doctoral Training Partnerships (training grant reference BB/T008741/1, studentships to LP and MB and BB/T008784/1 to EIF). EIF thanks the Ineos Oxford Institute for Antimicrobial Research for support. CLT, JS and AJM thank the Medical Research Council for support through the grant MR/T016035/1. CLT thanks the University of Bath Prize Fellowship Scheme and the UK Medical Research Council for fellowship funding (UKRI330).

