## Supplemental Figures and Tables for "Time-resolved ligand dynamics revealed in a β-lactamase using room-temperature serial crystallography"

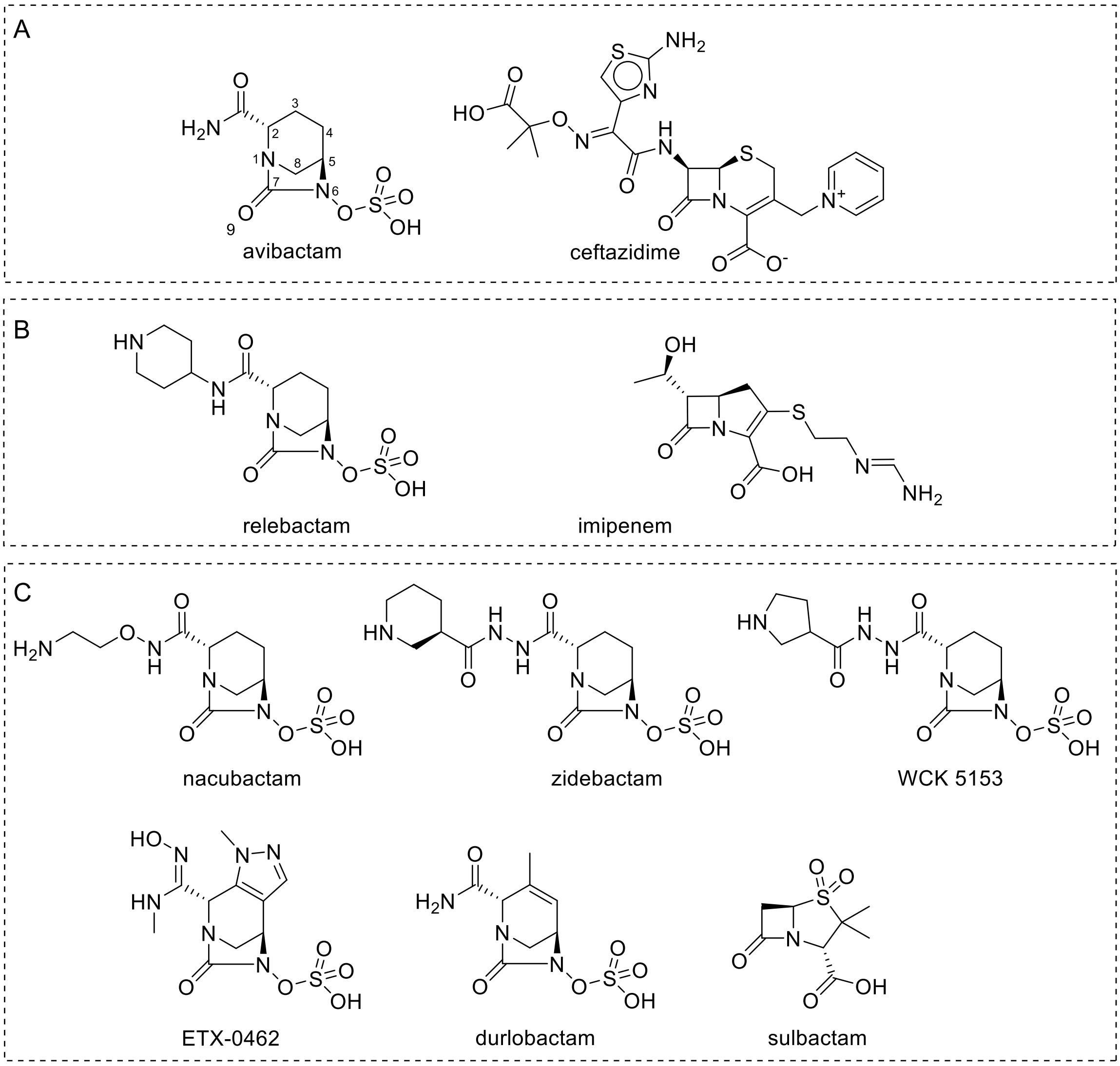


**Figure S1. Chemical structures of diazabicyclooctanes and partnered β-lactams. (A)** The clinically used Avicaz combination comprises avibactam (the first clinically available diazabicyclooctane (DBO)), left, with core atoms numbered, and ceftazidime, right, a cephalosporin antibiotic. **(B)** The clinically available DBO, relebactam, left, that is used in combination with the carbapenem β-lactam imipenem, right. **(C)** Examples of DBOs with evidenced dual action activity, targeting β-lactamases and penicillin binding proteins (PBPs). Durlobactam was approved for clinical use with its partner, the β-lactam β-lactamase inhibitor, sulbactam.

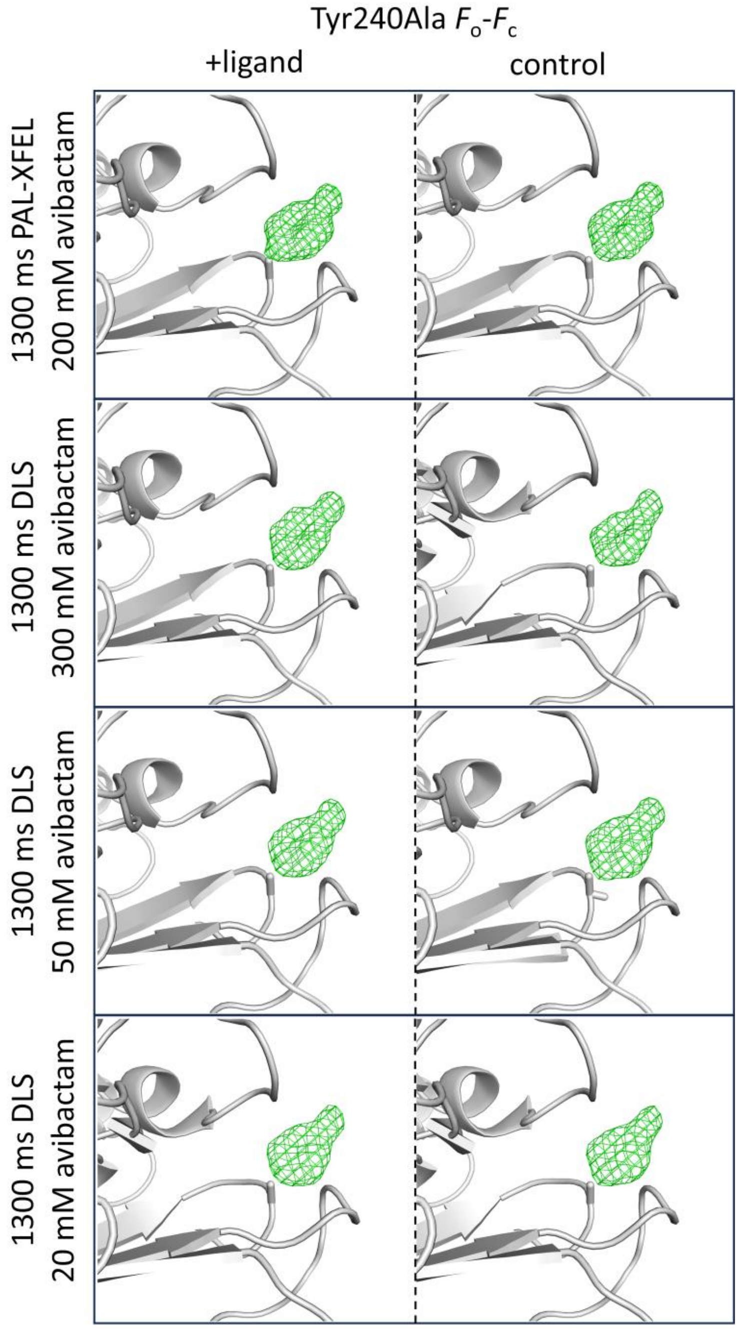


**Figure S2. Data quality verification with omit maps of Tyr240.** $F_o$-$F_c$ maps (green, contoured at 3.5σ), calculated after removal of Tyr240 side chain up to the Cβ atom (effectively a Tyr240Ala substitution), are shown for + ligand and control datasets for each drop on fixed target experiment (labelled on the *left*).

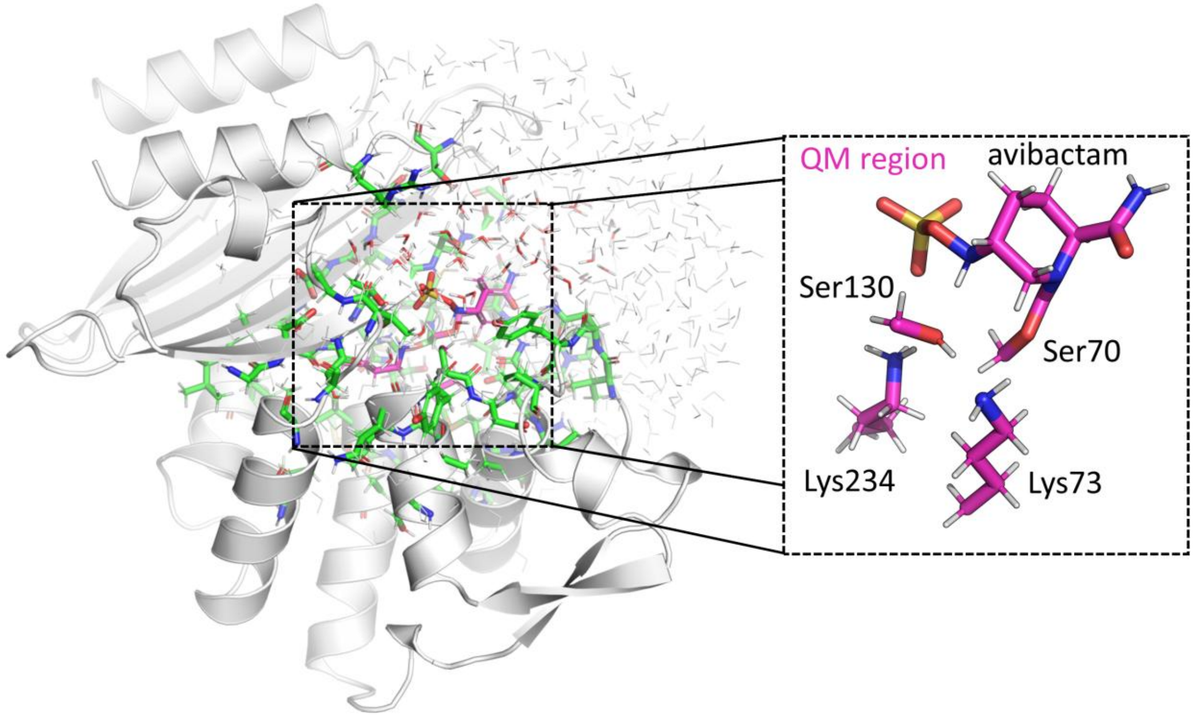


**Figure S3. Overall view of QM/MM geometry optimization calculation region.** Protein atoms fixed at the MM level during calculations are shown as grey cartoon, with waters as grey sticks. Residues and waters at the MM level which can have their atomic positions moved during optimisation are shown as sticks and coloured by atom with green as carbon, oxygen as red, nitrogen as blue and hydrogens white. These were picked as a result of being within 6 Å of the QM region (C atoms coloured in pink), comprising the carbamoyl avibactam-Ser70, Lys73, Lys234 and Ser130. A zoom in of the QM region is shown in the dashed box. Link atoms in the QM region are not shown.

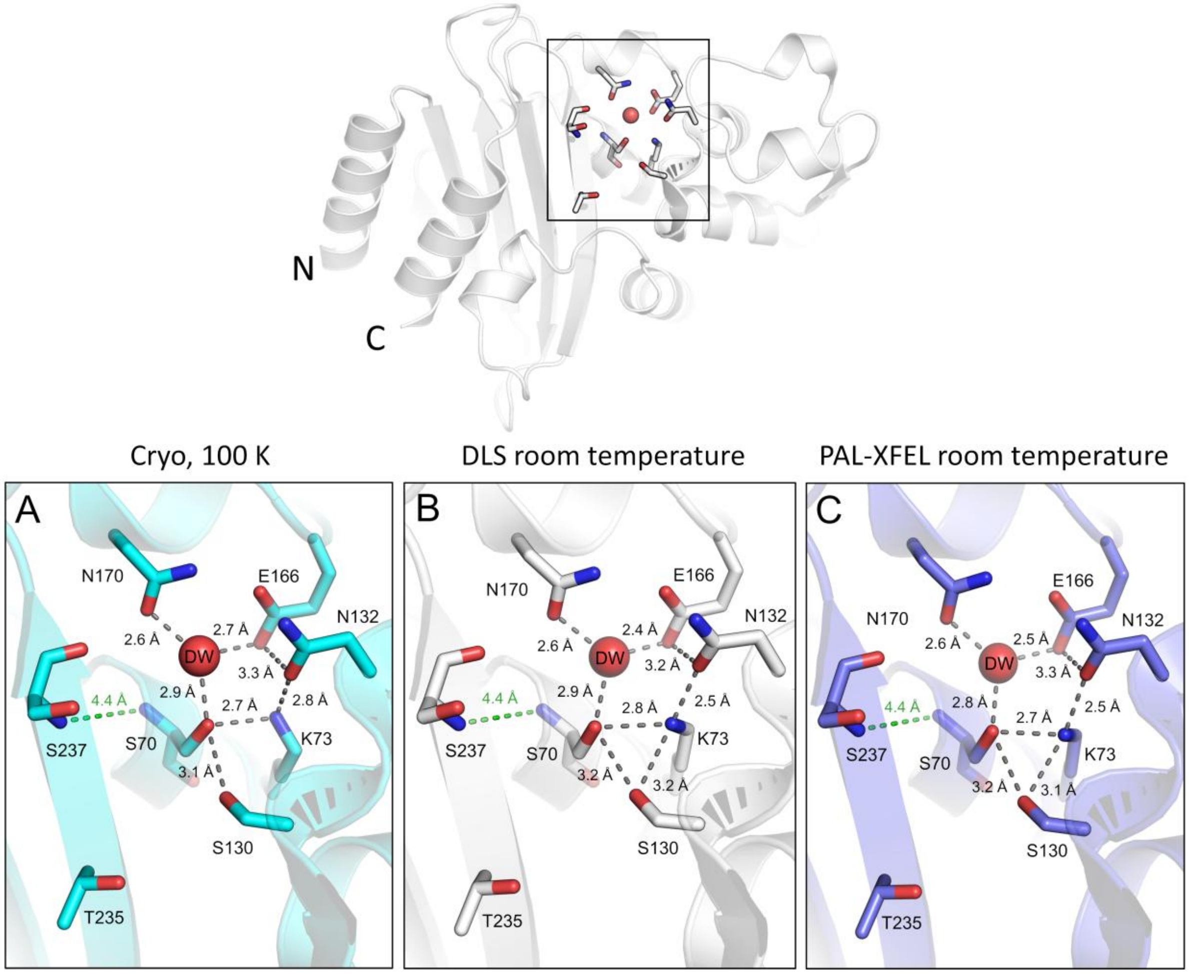


**Figure S4. Comparison of resting CTX-M-15 active sites from room temperature serial data and 100K cryo data.** *Top*, overall structure of CTX-M-15 (for clarity only the Diamond Light Source, DLS, room temperature structure is shown), with the active site boxed and N- and C-termini labelled. *Bottom***,** close up views of the active sites of **(A)** 100 K structure of apo CTX-M-15 (PDB 4HBT, 1.10 Å resolution), **(B)** Room temperature CTX-M-15 resting state data collected at DLS on the fixed target chips used in this study (1.73 Å resolution), **(C)** Room temperature CTX-M-15 resting state data collected at the X-ray Free Electron Laser facility PAL-XFEL (1.6 Å resolution). The hydrolytic water that is thought to be important for decarbamoylation (DBOs) or deacylation (β-lactams) of covalent complexes is labelled DW. Distances are labelled in Å for hydrogen bonds in the active site (grey dashes), and between backbone amides of Ser237 and Ser70 that constitute the oxyanion hole (green dashes). Crystals were grown in the same conditions for all three structures (0.1 M Tris pH 8.0, 2.0 -2.4 M $(NH_4)_2SO_4$). Note, there are small difference in the room temperature data, particularly, movement of Lys73 and a reduction in the distance between DW and Glu166.

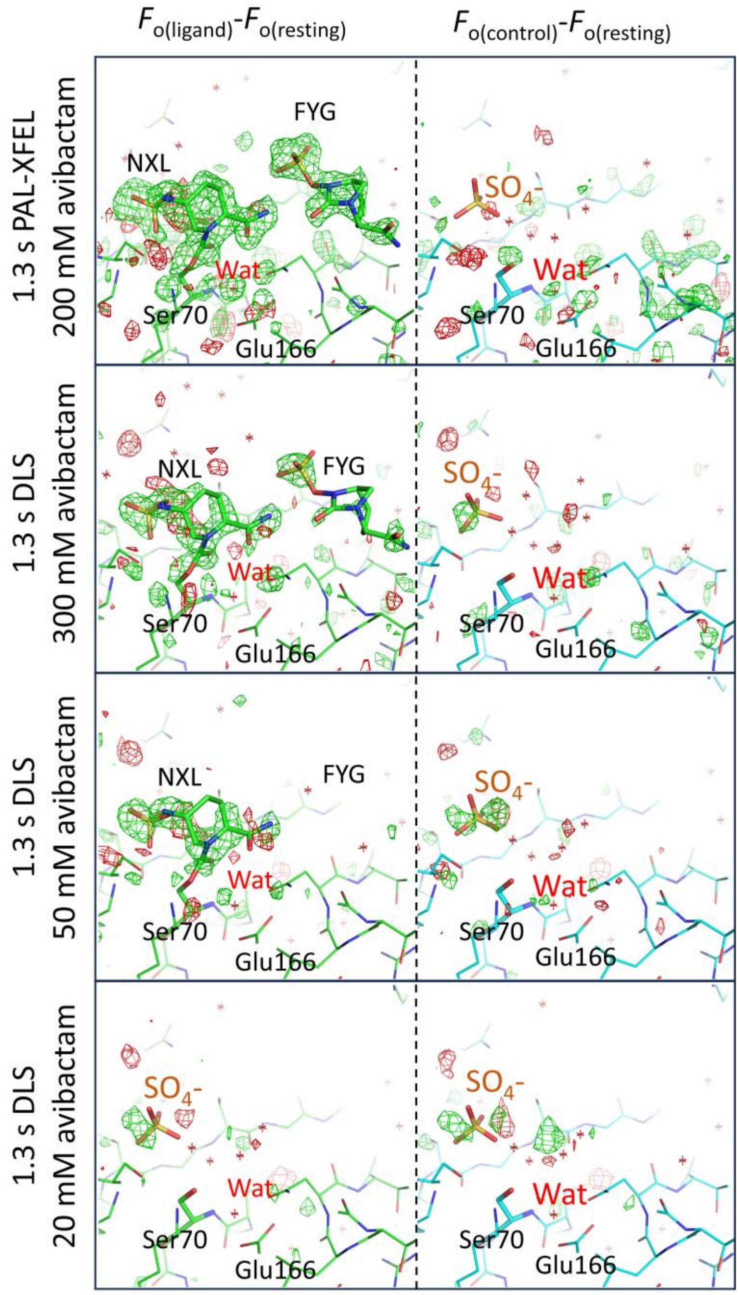


**Figure S5. Isomorphous difference maps of drop on fixed target mixing with room temperature resting state data.** Isomorphous difference maps for drop on fixed target mixing data (labelled on the left) and respective resting state data collected at each source (PAL-XFEL and DLS), contoured at +3σ (green) and -3σ (red). NXL is the covalently-bound, ring-opened avibactam and FYG the non-reacted avibactam. *Left*, isomorphous data from wells exposed to ligand against resting state data, $F_{o(ligand)}$-$F_{o(resting)}$; *right*, isomorphous data from wells not exposed to ligand against resting state data, $F_{o(control)}$-$F_{o(resting)}$.

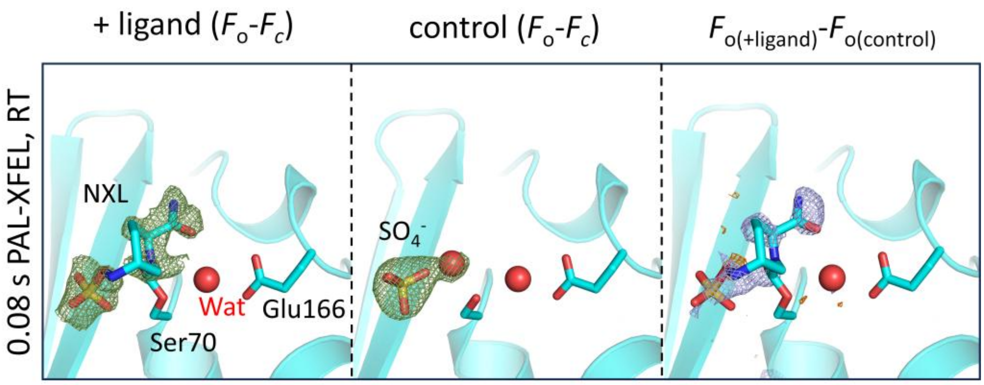


**Figure S6. Avibactam reaction with CTX-M-15 after 0.08 s.** Views from the active sites of avibactam bound in CTX-M-15 (NXL is the covalently bound avibactam) from room temperature serial crystallography data after 0.08 s (i.e. 80 ms) mixing using the drop on fixed target approach at PAL-XFEL. Electron densities (mesh) are contoured at 3σ around avibactam molecules or the active site sulphate and water. *Left*, $F_o$-$F_c$ electron density (green), from data collected from wells in which ligand was ejected (+ ligand, 1.5 Å resolution). *Middle*, $F_o$-$F_c$ electron density (green), in control wells in which ligand is not ejected (control, 1.55 Å resolution). *Right*, isomorphous difference maps ($F_o$-$F_o$) between +ligand and control, contoured at +(blue)/-(orange) 3σ. Note, we do not observe the non-covalent, intact avibactam (FYG) bound at the entrance to the active site.

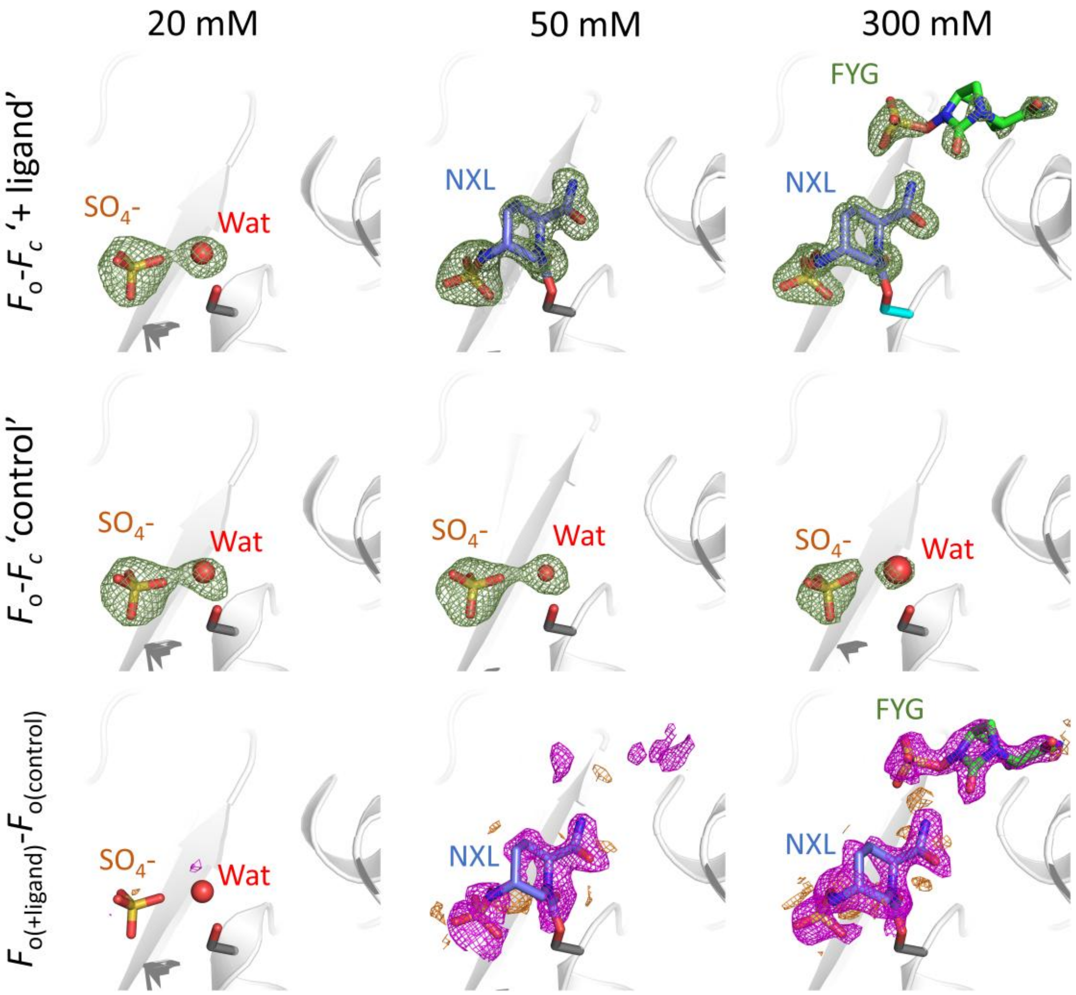


**Figure S7. Concentration dependence of avibactam binding at 1.3 s.** *Top*, $F_o$-$F_c$ OMIT electron density in the '+ ligand' active sites, contoured at 3 σ (green) around the $SO_4^-$ and active site water (Wat), or the covalently bound avibactam (purple C atoms, NXL) and/or intact avibactam at the opening to the active site (green C atoms, FYG). *Middle*, $F_o$-$F_c$ OMIT electron density in the 'control' active sites. Contoured as above. *Bottom*, $F_{o(+ligand)}$-$F_{o(control)}$ isomorphous difference maps contoured at 3 σ (+/-, purple/orange) within 10 Å of the active site 'Wat'.

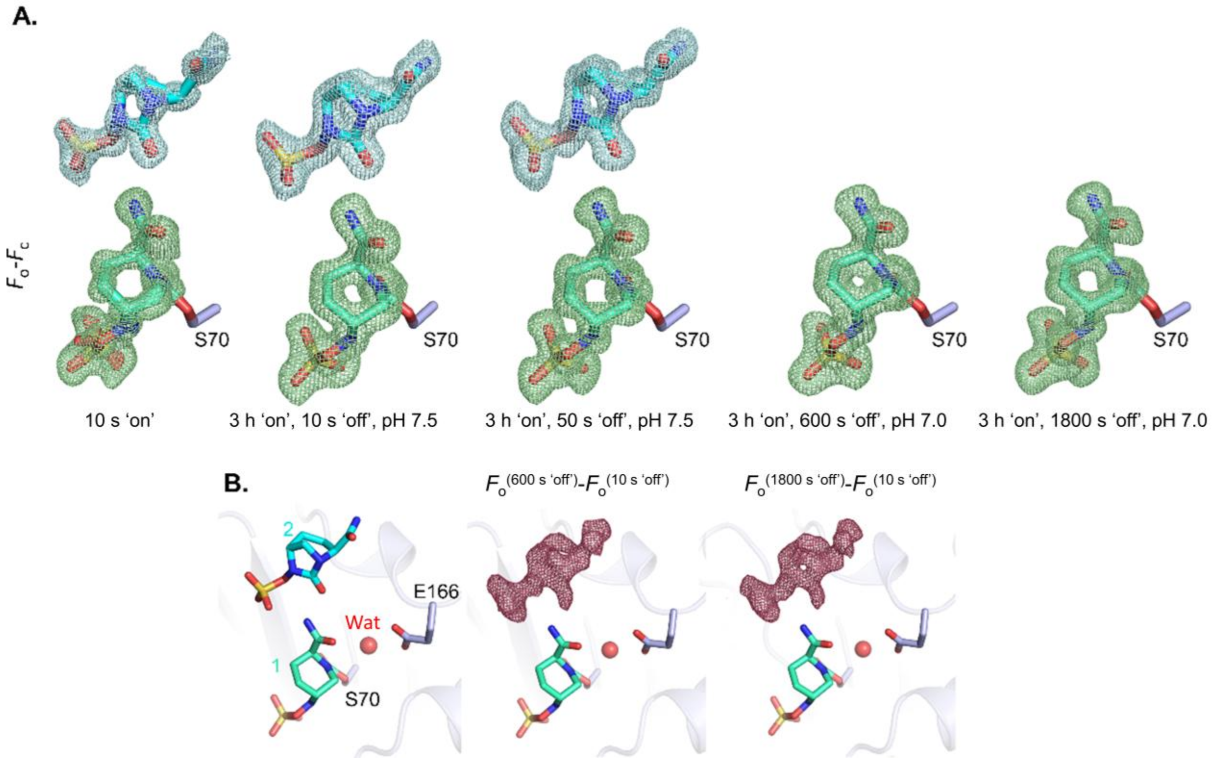


**Figure S8. Time dependence of avibactam soaking in macrocrystals from data collected at 100K.** Avibactam complexes were initially obtained by soaking complexes for 10 s (10 s 'on') or 3 h (3 h 'on'). Crystals were then washed for 10 s, 50 s, 600 s or 1800 s in crystallisation buffer without avibactam. See methods. **(A)** $F_o$-$F_c$ maps of covalent and non-covalent avibactam (3 σ, green and blue, respectively) within 10 s and a variety of soaks set up for 3hrs with several 'wash' times facilitated by a mild pH jump (pH 7.0-7.5) at 10 s, 50 s, 600 s, or 1800 s. The second, intact molecule of avibactam can be observed in a 10 s macrocrystal soak at 100 K. **(B)** This second molecule, but not the covalently reacted molecule, can be seen to leave after crystals were washed for 600 s or 1800 s, evidenced in $F_{o(timepoint)}$ -$F_{o(10s)}$ isomorphous difference maps, contoured at -3σ (red).

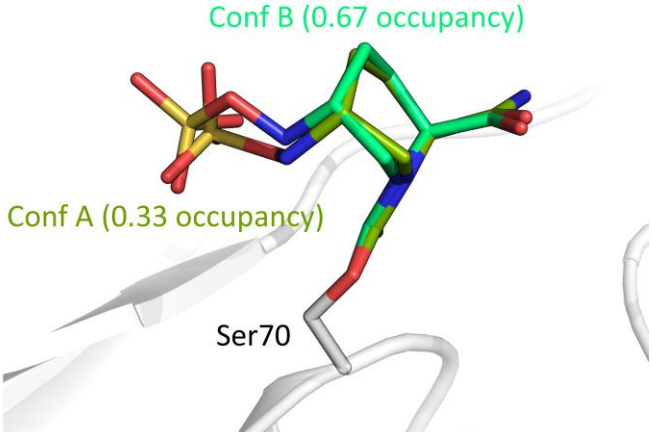


**Figure S9. Dual conformations of the covalent reacted avibactam conformations after 10 s exposure to avibactam.** View from the active site of the cryo-cooled structure of CTX-M-15 (0.94 Å resolution) after 10 s exposure of macro-crystals to avibactam. The covalently linked avibactam (NXL) is bound in two conformations, Conf A (green) and Conf B (lime), with occupancies of 0.33 and 0.67, respectively. Note, Conf B is in a similar conformation to that observed in the 600 s room temperature data and previous cryo-cooled 24-hour data (PDB 4HBU).

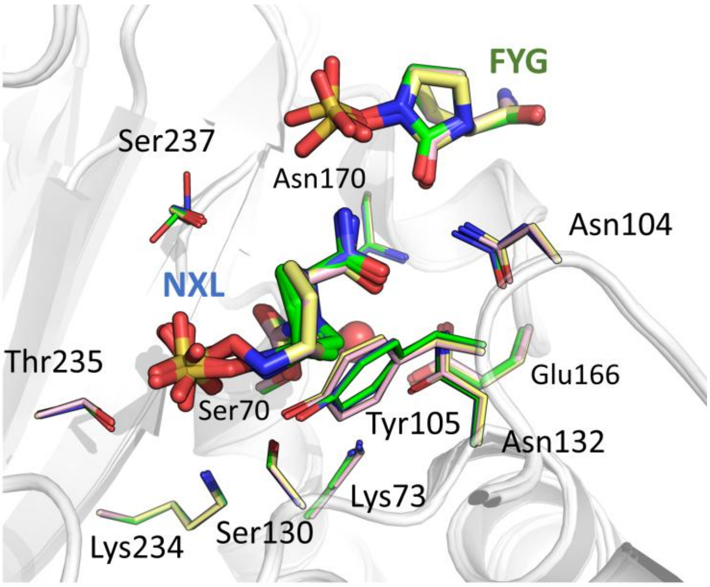


**Figure S10. Interactions in the CTX-M-15 active site with reacted avibactam.** (A) Overlay of the views from the active sites of avibactam bound in CTX-M-15 from 1.3 s (room temperature DLS, C atoms pink, and PAL-XFEL, C atoms yellow), 10 s (100K, C atoms green), 600 s (room temperature, C atoms cyan) and 24+ hours (100K, PDB 4HBU, C atoms blue) datasets. Residues that form hydrogen bonds with avibactam, interact hydrophobically or which are important for inhibition are shown as sticks. Details of the distances at each timepoint are shown in **Table S6**.

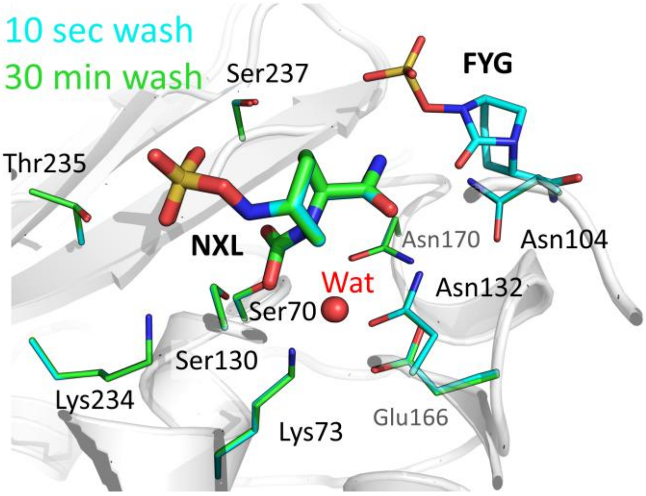


**Figure S11. Loss of intact avibactam from the CTX-M-15 active site does not affect binding of the covalently reacted avibactam.** Overlay of the active sites of CTX-M-15 soaked with avibactam for 3 hours, followed by soaking the crystal in buffer without avibactam, for 10 s (cyan C atoms), and for 1800 s (green C atoms).

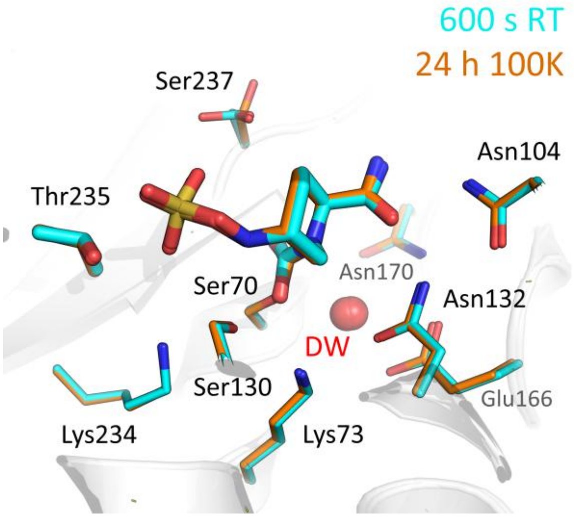


**Figure S12. Comparison of 600 s room temperature and 24-hour cryo-cooled data.** Superposed views from the active sites of the 600 s room temperature data and the previously determined co-crystal structure of avibactam with CTX-M-15. Note, both structures represent an 'in' conformation of the N6 N-sulfate nitrogen pointing towards the C7 of the carbonyl.

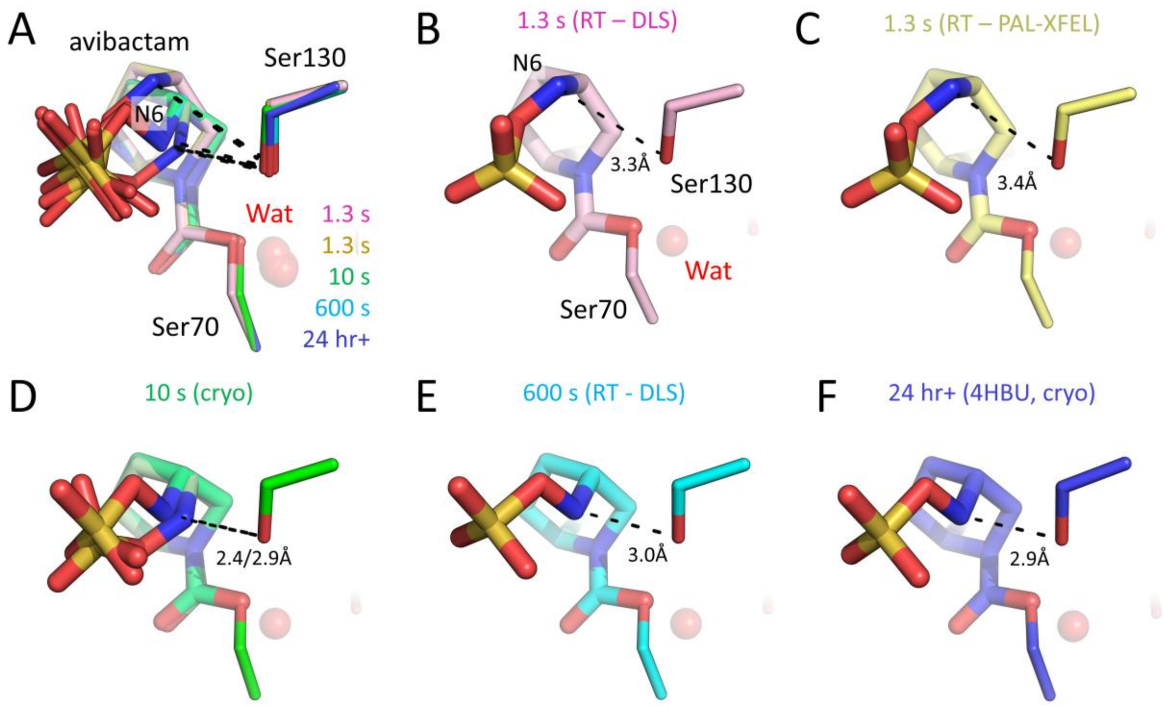


**Figure S13. Ser130 interactions with the N-sulfate nitrogen atom of avibactam at different timepoints.** Views from the active sites of avibactam covalently bound to Ser70 of CTX-M-15. The distances between Ser130 and the N6 of avibactam are shown as dashed lines, and the distance labelled in Å. **(A)** Overlay of CTX-M-15:avibactam structures, coloured according to time point as in main text figure 2. with the distance between N6 and Ser130 shown as dashed lines. **(B) – (F)** 1.3 s room temperature from DLS (pink) or PAL-XFEL (yellow), 10 sec 100 K (green, with avibactam in two conformations), 600 s room temperature from DLS (cyan) and previously published 100 K co-crystal structure (4HBU, blue).

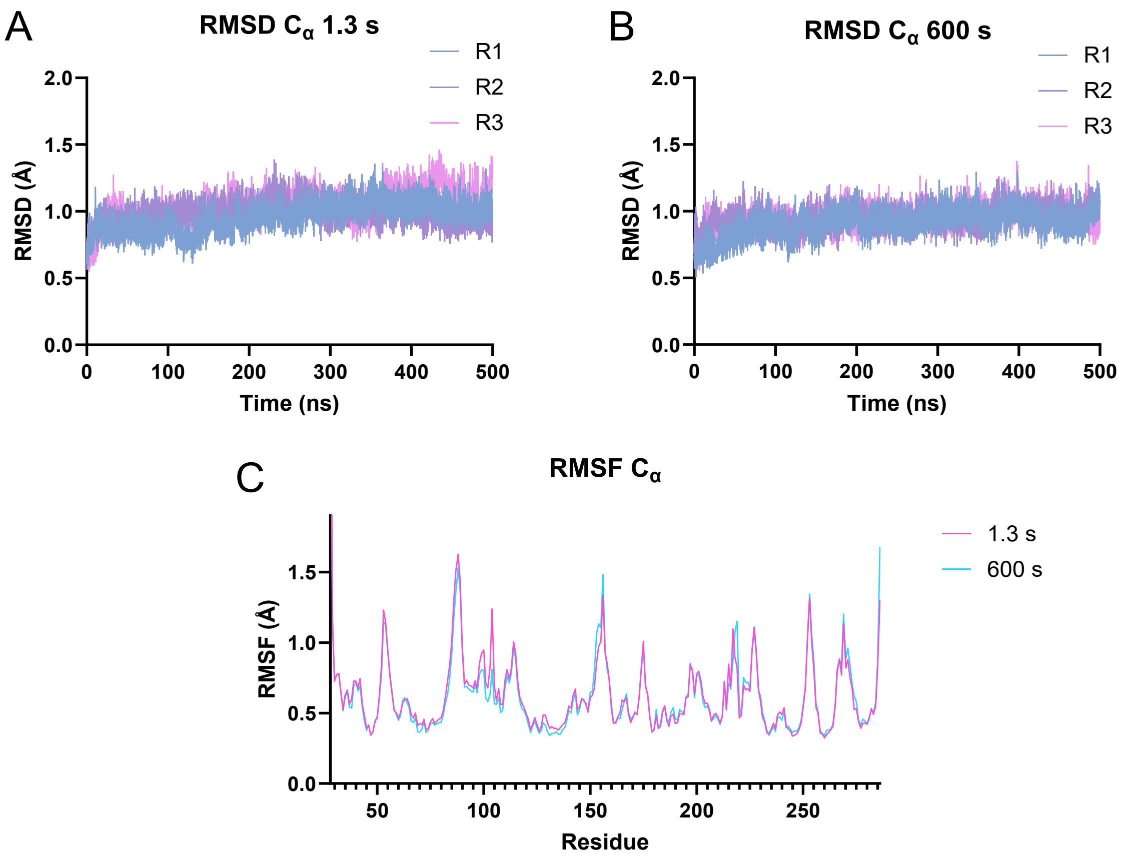


**Figure S14. Molecular dynamics trajectories.** $C_\alpha$ root mean square deviations (RMSDs, in Å) over the course of the three 500 ns repeat trajectories (R1 – R3) of the room temperature **(A)** 1.3 s PAL-FEL structure and **(B)** 600 s DLS structure. **(C)** $C_\alpha$ root mean square fluctuations (RMSFs) averaged over the three repeats for both the 1.3 s and 600 s structures.

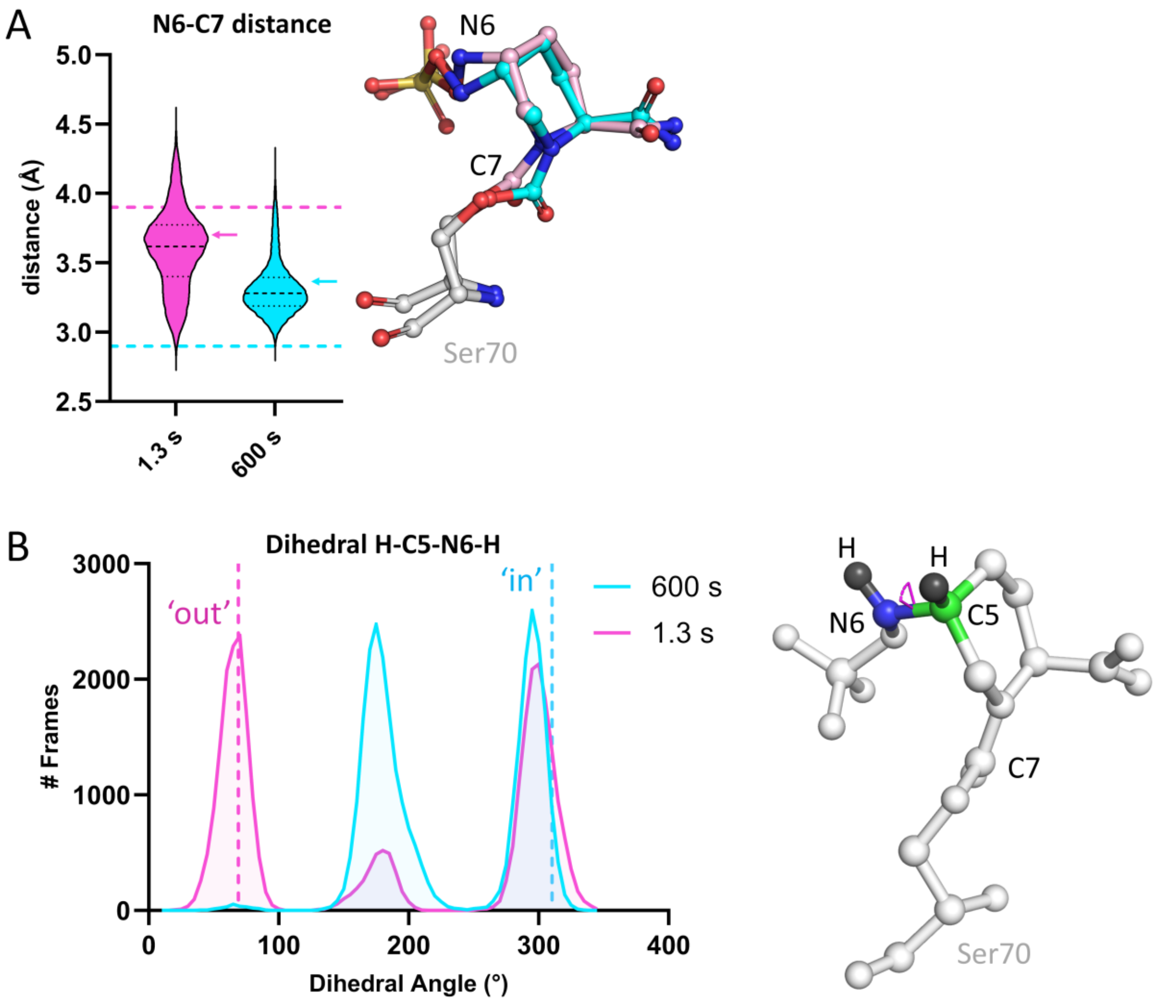


**Figure S15. Dynamics of the *N*-sulfate nitrogen with respect to C7 in the avibactam carbamoyl complex.** Analysis of the 3 x 500 ns molecular dynamics simulations, starting from the 1.3 s X-ray structure (pink) and 600 s structure (blue). Pink and blue dashed lines indicate the distance in the respective X-ray crystal structures and the 'in' and 'out' conformations. **(A)** *Left*, Violin plots of the distribution of the distance between N6 and C7 over the course of the simulations. Minor black dashed lines represent the quartiles, and the major dashed line the median of each dataset. *Right*, snapshots of representative frames at the median distance (indicated by coloured arrows in the graph) in each simulation indicating the 'out' and 'in' AVI conformations, 1.3 s (pink) and 600 s (blue), respectively. **(B)** *Left*, Frequency distribution of the H-C5-N6-H dihedral angle, binned every 5° between 0 and 360°. Dashed lines show the angles in the respective crystal structures ('out', 1.3 s; 'in' 600 s). *Right*, the H-C5-N6-H dihedral angle (pink) is highlighted. Example shown is from the 1.3 s PAL-XFEL structure.

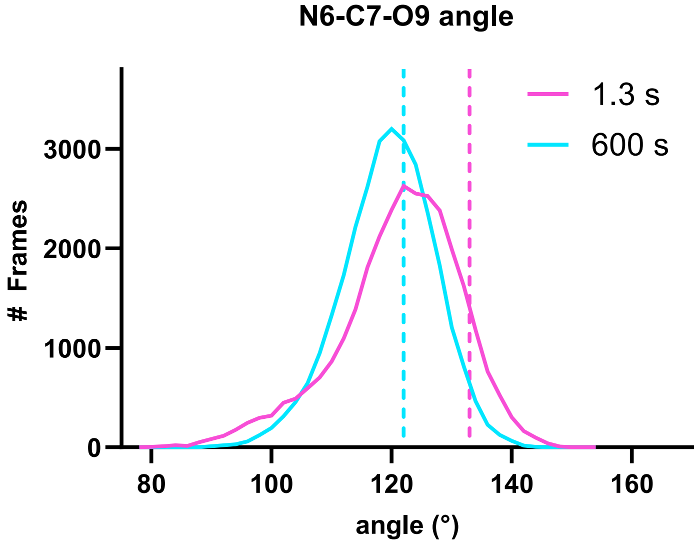


**Figure S16. Distribution of the N6-C7-O9 angle over the molecular dynamics simulations.** Frequency distribution (binned every 2°) of the measurement of the N6-C7-O9 angle (also known as the Bürgi-Dunitz angle ($\alpha_{BD}$) – see Discussion) over 3 x 500 ns molecular dynamics simulations starting from 1.3 s (pink) and 600 s (blue) room temperature crystal structures. Dashed lines show the angles in the 1.3 s (133°, pink) and 600 s (122°, blue) crystal structures.

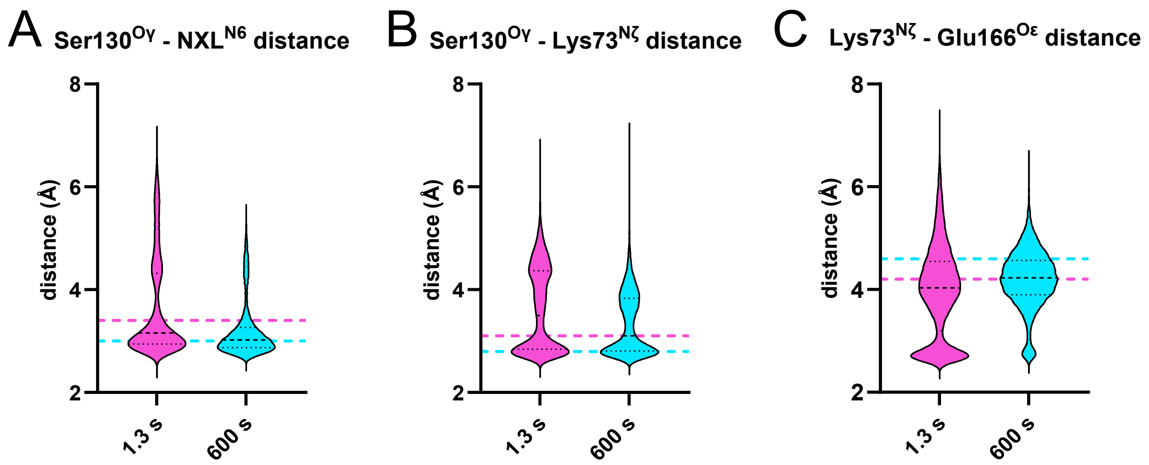


**Figure S17. Distances between key residues and the avibactam carbamoyl complex over the molecular dynamics simulations.** Violin plots showing the distribution of distances for 1.5 µs simulations (pink, 1.3 s; cyan, 600 s), for **(A)** Ser130$^{O\gamma}$ – NXL$^{N6}$; **(B)** Ser130$^{O\gamma}$ – Lys73$^{N\zeta}$; **(C)** Lys73$^{N\zeta}$ – Glu166$^{O\varepsilon}$. Dashed lines show distances in the respective crystal structures.

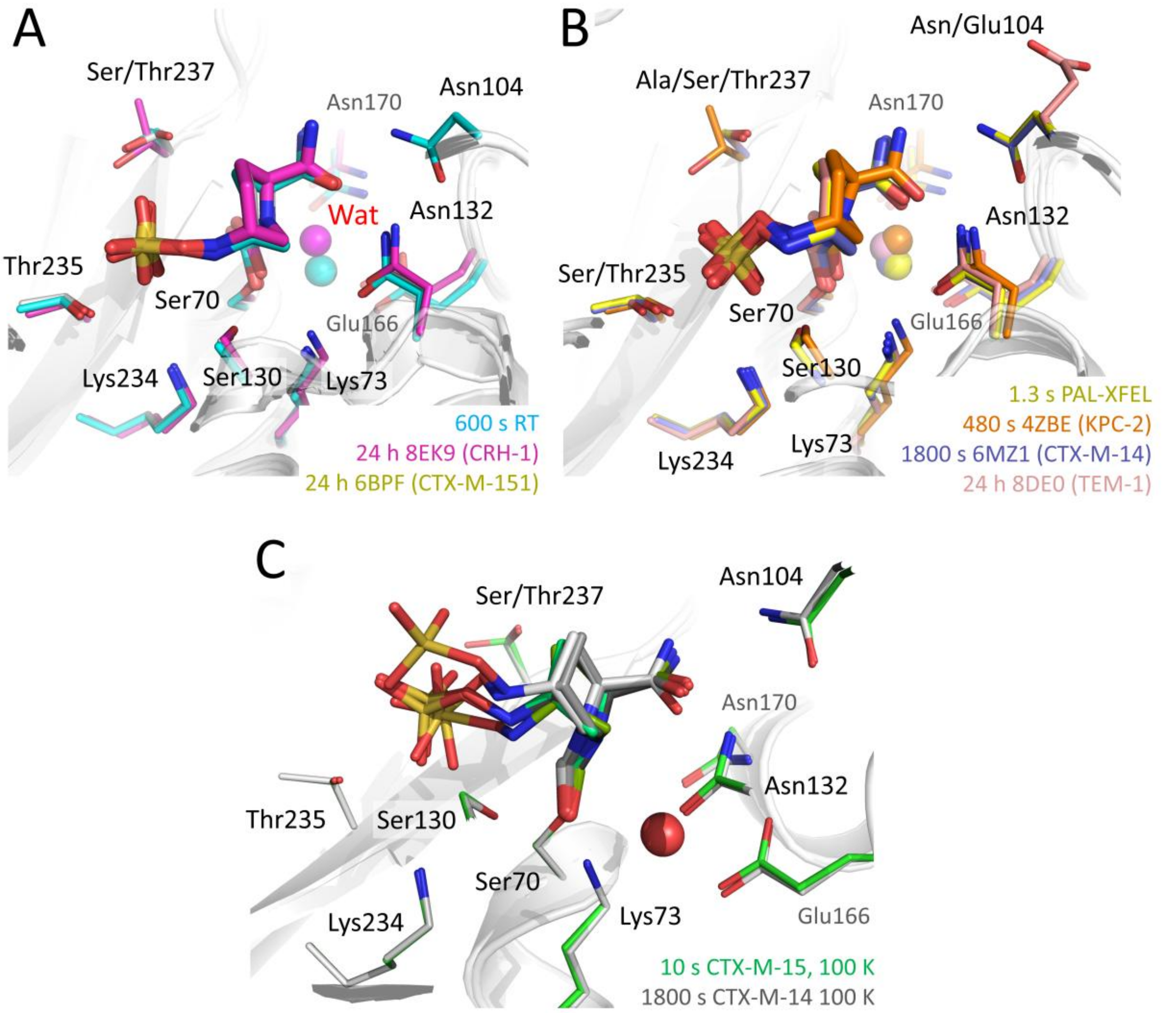


**Figure S18. Comparisons of avibactam binding in serine-β-lactamases.** Close up views from the active sites of selected previously published Ser70-avibactam carbamoyl complexes compared with structures presented here. See sequence alignment in **Figure S19**. **(A)** Comparison of our CTX-M-15 room temperature 600 s structure (cyan, 1.7 Å resolution, PDB 9TOH) with CRH-1 (PDB 8EK9, pink, 1.4 Å resolution, 24-hour soak, RMSD over 261 Cα with CTX-M-15 1.04 Å) and CTX-M-151 (PDB 6BPF, yellow, 1.32 Å resolution, 24-hour soak, RMSD over 262 Cα with CTX-M-15 0.63 Å), showing the 'in' conformation for the N-sulfate moiety of avibactam. **(B)** Comparison of our CTX-M-15 room temperature 1.3 s structure collected at PAL-XFEL (yellow, 1.45 Å, PDB 9TO3) with KPC-2 (PDB 4ZBE, orange, 1.8 Å resolution, 8 min soak, RMSD over 259 Cα atoms with CTX-M-15 1.31 Å), CTX-M-14 (PDB 6MZ1, yellow, 1.0 Å resolution, 30 min soak, RMSD over 261 Cα with CTX-M-15 0.71 Å) and TEM-1 (PDB 8DE0, 1.72 Å resolution, 24 hour soak, RMSD over 252 Cα with CTX-M-15 1.76 Å) showing the 'out' conformation for the *N*-sulfate moiety of avibactam. **(C)** Conformations A and B in the 10 sec CTX-M-15 structure (this work, light and dark green, PDB 9TOC) and 1800 s CTX-M-14 (light and dark grey, PDB 6MZ2, 0.83 Å).

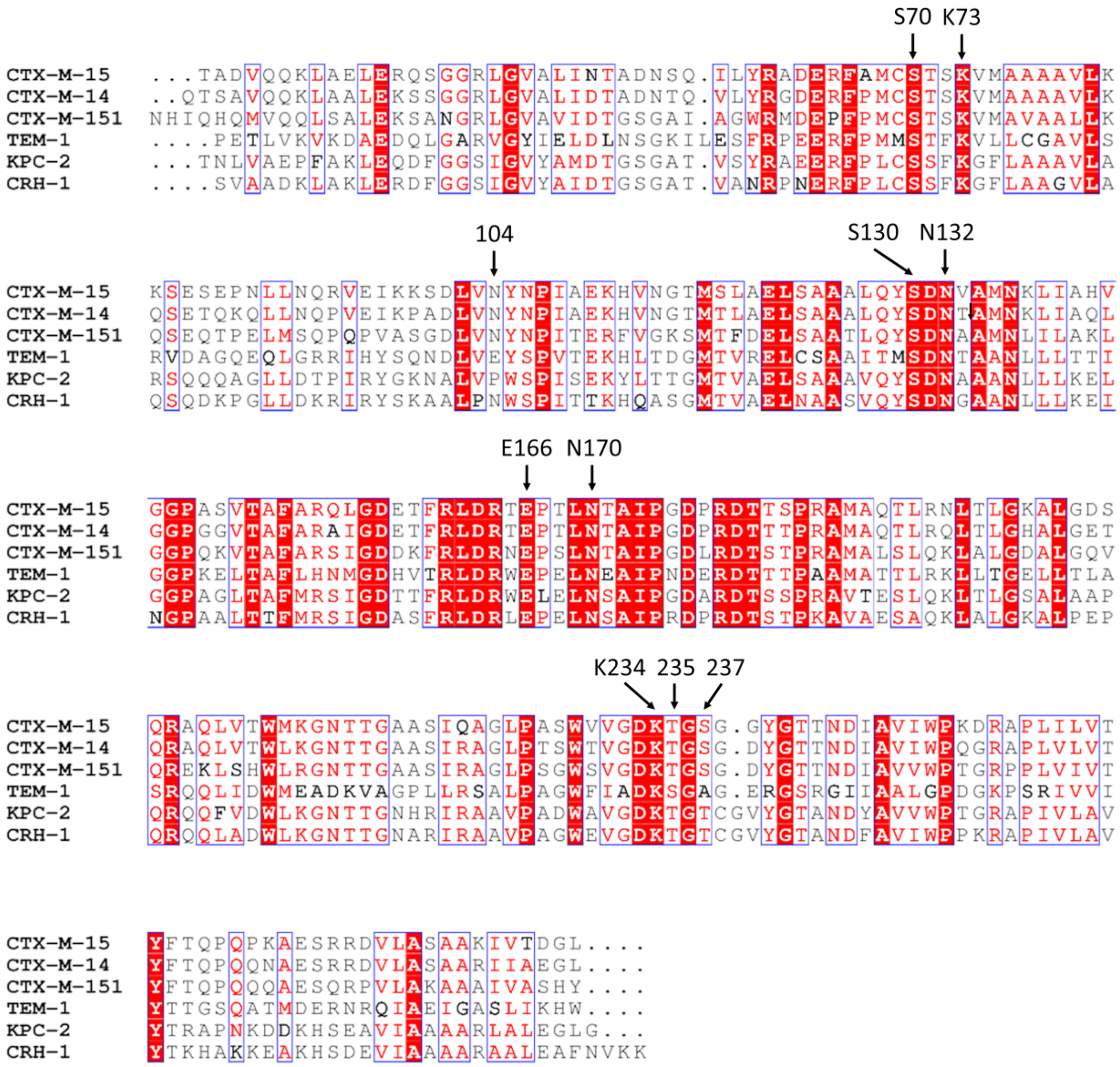


**Figure S19. Sequence alignment of selected class A serine-β-lactamases.** Aligned sequences of CTX-M-15, CTX-M-14, CTX-M-151, TEM-1, KPC-2 and CRH-1 from PDB structures in **Figure S18**. Some key active site residues (those shown in **Figure S18** that are involved in avibactam binding) are labelled. Strictly conserved residues are boxed in white on a red background and highly conserved residues are boxed in red on a white background. Sequences were aligned with Clustal Omega (1), and the figure generated with ESPript 3.0 (2).

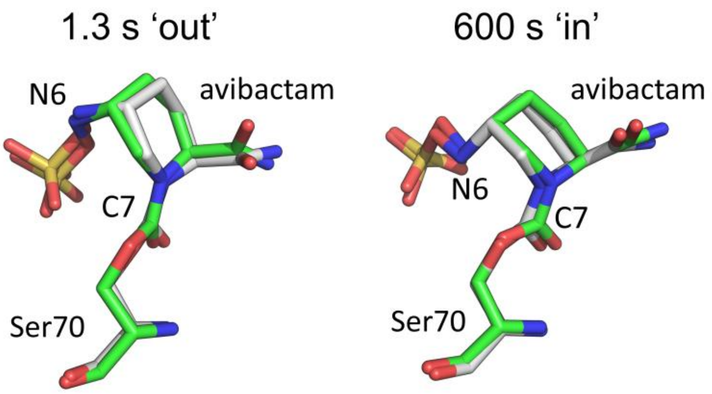


**Figure S20. Molecular dynamics equilibrated structures.** Close up views of the Ser70-avibactam carbamoyl complexes. The equilibrated structures for molecular dynamics (MD) simulations are shown in green, overlaid with the crystal structures (gray) that were used as the starting structures for the respective MM MD simulations. N6 and C7 atoms are labelled. *Left*, structures from the PAL-XFEL 1.3 s drop on fixed target room temperature timepoint are in the 'out' conformation, in which N6 is pointing away from C7. *Right*, structures from the 600 s pre-mixed room temperature data are in the 'in' conformation, in which N6 is pointing towards C7.

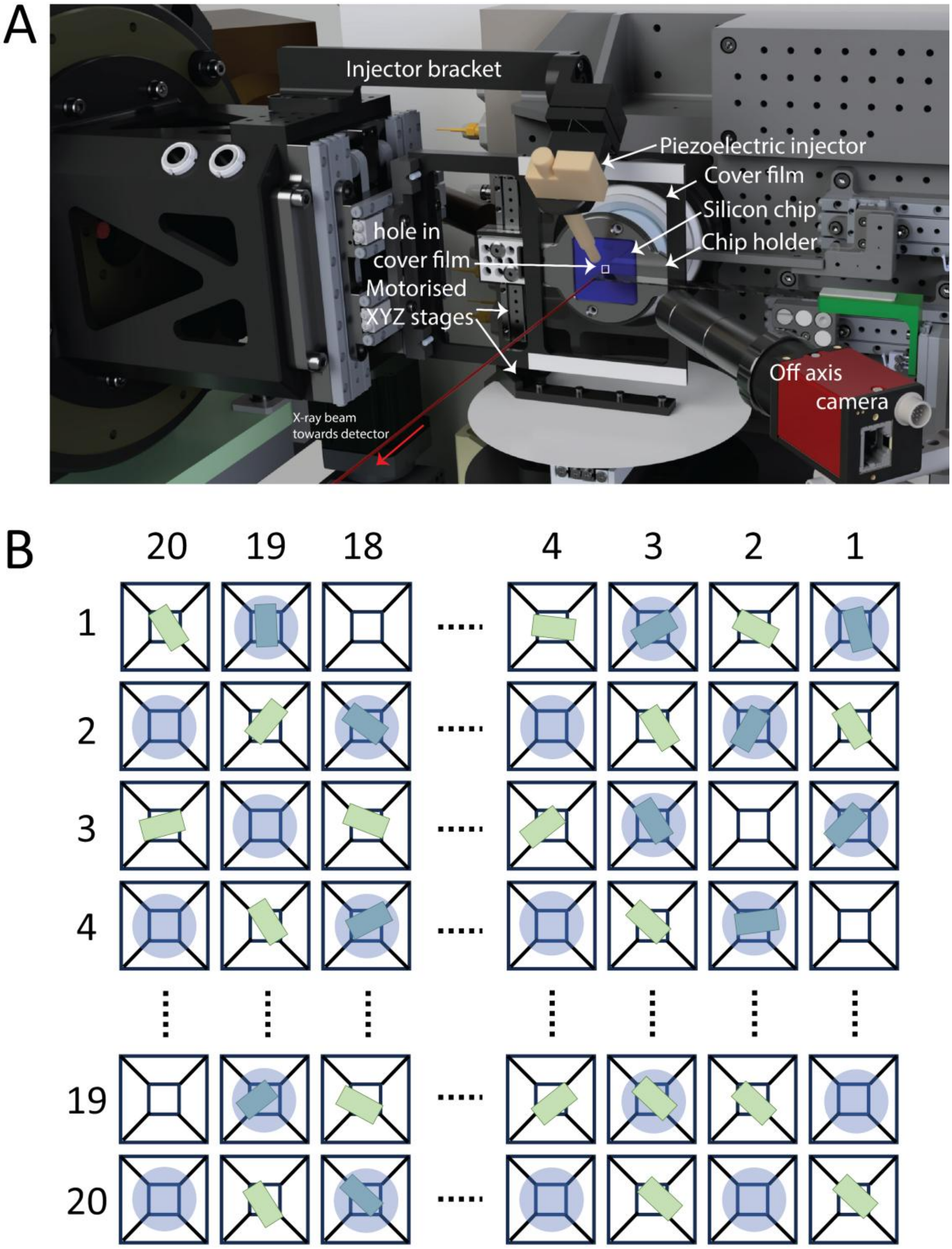


**Figure S21. Drop on fixed target setup and checkerboard control pattern. (A)** The drop on fixed target as setup at PAL-XFEL and DLS. The piezoelectric injector (PEI) adds ligands to the silicon chip (containing an open face) that is covered by a cover film with a hole for the PEI. An off axis camera is used to align droplets with the wells. **(B)** A schematic of the checkerboard pattern of droplet addition by the PEI. Crystals (green rectangles) sit in the wells of the chip. Droplets are added to every other well, and every well is exposed to X-rays with subsequent data split into '+ligand' and 'control' datasets.

**Table S1. Room temperature PAL-XFEL and DLS crystallographic data collection and refinement statistics for CTX-M-15 crystals after 1.3 s avibactam exposure**

| **Beamline** | PAL-XFEL NCI - SFX | | | DLS I24 | | |
|---|---|---|---|---|---|---|
| **Timepoint** | resting | 1.3 s | | resting | 1.3 s | |
| **+ ligand / control** | N/A | + ligand | control | N/A | + ligand | control |
| **PDB accession** | 9TO1 | 9TO3 | 9TO4 | 9TO5 | 9TO6 | 9TO7 |
| ***Data collection*** | | | | | | |
| Space group | $P2_12_12_1$ | $P2_12_12_1$ | $P2_12_12_1$ | $P2_12_12_1$ | $P2_12_12_1$ | $P2_12_12_1$ |
| Molecules/ASU | 1 | 1 | 1 | 1 | 1 | 1 |
| Cell dimensions | | | | | | |
| *a*, *b*, *c* (Å) | 45.29, 45.94, 118.45 | 45.26, 45.98, 118.88 | 45.27, 45.88, 118.65 | 45.19, 45.69, 118.36 | 45.23, 45.85, 118.68 | 45.25, 45.75, 118.47 |
| α=β=γ (°) | 90 | 90 | 90 | 90 | 90 | 90 |
| # 'chips' | 1 | 2 | 2 | 2 | 4 | 4 |
| # integrated lattices | 16444 | 13377 | 14176 | 10756 | 10368 | 9773 |
| Wavelength (Å) | 1.30419 | 1.304197 | 1.304197 | 0.9998 | 0.6199 | 0.6199 |
| Resolution (Å) | 59.23 – 1.6 (1.6276 – 1.6) | 59.44 – 1.45 (1.475 – 1.45) | 59.32 – 1.47 (1.4953 – 1.47) | 59.18 – 1.73 (1.7598 – 1.73) | 59.34 – 1.62 (1.679 – 1.62) | 59.24 – 1.62 (1.679 – 1.62) |
| $R_{split}$ | 14.9 (38.3) | 18.5 (71.9) | 19.5 (89.2) | 17.4 (45.3) | 20.2 (43.9) | 23.7 (49.4) |
| CC 1/2 | 96.9 (68.9) | 95.6 (34.8) | 95.6 (38.1) | 95.6 (59.4) | 94.2 (31.4) | 91.2 (30.8) |
| $I / \sigma I$ | 4.286 (1.044) | 2.996 (0.612) | 3.043 (0.654) | 3.547 (1.341) | 3.237 (1.413) | 2.922 (1.328) |
| Completeness (%) | 99.99 (99.94) | 100 (100) | 100 (100) | 99.95 (99.38) | 99.99 (100) | 99.99 (99.94) |
| Multiplicity | 78.90 (11.77) | 65.99 (15.49) | 63.01 (17.33) | 34.42 (9.18) | 43.71 (17.54) | 37.49 (15.06) |
| ***Refinement*** | | | | | | |
| Resolution (Å) | 42.83 – 1.60 | 42.88 – 1.45 | 42.79 – 1.47 | 59.18 – 1.73 | 59.34 – 1.62 | 59.24 – 1.62 |
| No. reflections | 33453 | 44760 | 42833 | 26353 | 32185 | 32068 |
| $R_{work}$ / $R_{free}$ | 13.98/16.48 | 19.07/14.95 | 19.33/15.70 | 15.79/18.86 | 22.99/18.49 | 25.15/20.55 |
| #. non-H atoms | | | | | | |
| Protein | 2012 | 2074 | 2012 | 1976 | 2080 | 1984 |
| Solvent | 218 | 188 | 180 | 193 | 247 | 230 |
| Ligand – NXL | N/A | 17 | N/A | N/A | 17 | N/A |
| Ligand – FYG | N/A | 17 | N/A | N/A | 17 | N/A |
| B-factors ($Å^2$) | | | | | | |
| Protein | 18.5 | 21.0 | 22.3 | 20.9 | 16.8 | 16.1 |
| Solvent | 32.7 | 37.6 | 38.4 | 33.2 | 31.5 | 30.1 |
| Ligand – NXL | N/A | 23.8 | N/A | N/A | 18.7 | N/A |
| Ligand – FYG | N/A | 33.2 | N/A | N/A | 22.7 | N/A |
| R.m.s. deviations | | | | | | |
| Bond lengths (Å) | 0.008 | 0.005 | 0.004 | 0.009 | 0.006 | 0.006 |
| Bond angles (°) | 0.994 | 0.756 | 0.724 | 1.044 | 0.823 | 0.752 |
| Ramachandran (%) | | | | | | |
| Outliers | 0.39 | 0.39 | 0.39 | 0.39 | 0.39 | 0.39 |
| Favoured | 98.44 | 98.46 | 98.44 | 98.44 | 98.46 | 98.44 |

Values in parenthesis are for the high-resolution shell; NXL – covalently reacted avibactam; FYG – intact non-covalently bound avibactam

**Table S2. Occupancies and PDB RSCCs for ligands bound in CTX-M-15 crystal structures.**

| Structure | NXL occupancy | NXL RSCC | NXL adjusted B-factor[#] | FYG occupancy | FYG RSCC | FYG adjusted B-factor[#] |
|---|---|---|---|---|---|---|
| PAL-FEL 1.3 s '+200 mM avibactam' | 1.0 | 0.96 | 0.94 | 0.66 | 0.91 | 1.29 |
| DLS 1.3 s '+300 mM avibactam' | 1.0 | 1.00 | 1.11 | 0.62 | 0.99 | 1.35 |
| DLS 1.3 s '+50 mM avibactam' | 0.78 | 0.94 | 1.17 | N/A | N/A | N/A |
| DLS 600 s | 1.00 | 0.98 | 0.71 | 0.76 | 0.96 | 1.36 |
| cryo 10 s | 0.33 (A)<br>0.67 (B*) | 0.97 (A)<br>0.97 (B*) | 1.97 (A)<br>1.27 (B*) | 0.64 | 0.93 | 1.77 |
| cryo 10 s 'off' | 1.00 | 0.99 | 0.94 | 0.86 | 0.96 | 1.29 |
| cryo 50 s 'wash' | 1.00 | 0.99 | 0.92 | 0.87 | 0.97 | 1.41 |
| cryo 600 s 'wash' | 1.00 | 0.98 | 1.04 | N/A | N/A | N/A |
| cryo 1800 s min 'wash' | 1.00 | 0.98 | 0.99 | N/A | N/A | N/A |

[#]ligand B-factor/protein B-factor

*Note, conf B is the same conformation as that observed in the 600 s room temperature structure and other cryo datasets.

NXL – covalently reacted avibactam; FYG – intact non-covalently bound avibactam

**Table S3. Room temperature serial crystallographic data collection and refinement statistics for CTX-M-15 crystals mixed with varying concentrations of avibactam**

| **Timepoint** | 1.3 s | | | |
|---|---|---|---|---|
| **Ligand concentration#** | 50 mM | | 20 mM | |
| **+ ligand / control** | + ligand | control | + ligand | control |
| **PDB accession** | 9TO8 | 9TO9 | 9TOA | 9TOB |
| ***Data collection*** | | | | |
| Beamline | DLS I24 | DLS I24 | DLS I24 | DLS I24 |
| Space group | $P2_12_12_1$ | $P2_12_12_1$ | $P2_12_12_1$ | $P2_12_12_1$ |
| Molecules/ASU | 1 | 1 | 1 | 1 |
| Cell dimensions | | | | |
| *a*, *b*, *c* (Å) | 45.22, 45.85, 118.75 | 45.22, 45.78, 118.61 | 45.22, 45.80, 118.64 | 45.22, 45.80, 118.67 |
| α=β=γ (°) | 90 | 90 | 90 | 90 |
| #chips | 3 | 3 | 3 | 3 |
| #indexed lattices | 11123 | 11061 | 9458 | 9694 |
| Wavelength (Å) | 0.9999 | 0.9999 | 0.9999 | 0.9999 |
| Resolution (Å) | 59.375 – 1.759 (1.789 – 1.759) | 59.306 – 1.736 (1.766– 1.736) | 59.322 – 1.92 (1.953– 1.92) | 59.333 – 1.92 (1.953– 1.92) |
| $R_{split}$ | 14.8 (51.1) | 16.1 (57.6) | 16.1 (40.1) | 15.9 (40.5) |
| CC 1/2 | 97.5 (44.8) | 96.8 (45.3) | 96.8 (61.5) | 96.7 (35.8) |
| $I / \sigma I$ | 4.144 (1.450) | 3.744 (1.075) | 4.153 (1.975) | 4.304 (1.992) |
| Completeness (%) | 100.0 (100.0) | 100.0 (100.0) | 100.0 (100.0) | 100.0 (100.0) |
| Multiplicity | 65.97 (37.14) | 61.09 (31.78) | 66.0 (38.91) | 67.95 (40.38) |
| ***Refinement*** | | | | |
| Resolution (Å) | 59.38 – 1.76 | 59.31 – 1.74 | 59.32 – 1.92 | 59.33 – 1.92 |
| No. reflections | 25282 | 26190 | 19525 | 19522 |
| $R_{work}$ / $R_{free}$ | 20.02/15.85 | 19.78/15.9 | 20.58/15.96 | 19.77/16.16 |
| #. non-H atoms | | | | |
| Protein | 2084 | 1995 | 1995 | 1994 |
| Solvent | 173 | 161 | 185 | 177 |
| Ligand – NXL | 17 | N/A | N/A | N/A |
| Ligand – FYG | N/A | N/A | N/A | N/A |
| B-factors (Å$^2$) | | | | |
| Protein | 23.7 | 25.6 | 25.4 | 25.3 |
| Solvent | 35.2 | 37.3 | 37.9 | 36.4 |
| Ligand – NXL | 27.8 | N/A | N/A | N/A |
| Ligand – FYG | N/A | N/A | N/A | N/A |
| R.m.s. deviations | | | | |
| Bond lengths (Å) | 0.005 | 0.009 | 0.006 | 0.006 |
| Bond angles (°) | 0.773 | 1.017 | 0.761 | 0.763 |
| Ramachandran (%) | | | | |
| Outliers | 0.39 | 0.39 | 0.39 | 0.39 |
| Favoured | 98.46 | 98.44 | 98.44 | 98.44 |

#Concentration of ligand in injector; Values in parenthesis are for the high-resolution shell; NXL – covalently reacted avibactam; FYG – intact non-covalently bound avibactam

**Table S4. Cryo-cooled crystallographic data collection at 100 K and refinement statistics for CTX-M-15 crystals mixed with avibactam**

| **Time point** | 10 s 'soak' | 'wash' 10 s | 'wash' 50 s | 'wash' 600 s (pH 7) | 'wash' 1800 s |
|---|---|---|---|---|---|
| **PDB accession** | 9TOC | 9TOD | 9TOE | 9TOF | 9TOG |
| ***Data collection*** | | | | | |
| Beamline | DLS I03 | DLS I03 | DLS I03 | DLS I03 | DLS I03 |
| Space group | $P2_12_12_1$ | $P2_12_12_1$ | $P2_12_12_1$ | $P2_12_12_1$ | $P2_12_12_1$ |
| Molecules/ASU | 1 | 1 | 1 | 1 | 1 |
| Cell dimensions | | | | | |
| *a*, *b*, *c* (Å) | 44.42, 45.59, 117.68 | 44.50, 45.60, 117.93 | 44.54, 45.64, 117.91 | 44.43, 45.59, 117.66 | 44.47, 45.57, 117.70 |
| α=β=γ (°) | 90 | 90 | 90 | 90 | 90 |
| Wavelength (Å) | 0.69998 | 0.69998 | 0.69998 | 0.69998 | 0.69998 |
| Resolution (Å) | 44.42 – 0.94 (0.96 – 0.94) | 44.5 – 1.22 (1.24 – 1.22) | 44.54 – 1.01 (1.03 – 1.01) | 44.43 – 1.24 (1.26 – 1.24) | 44.47 – 0.87 (0.89 – 0.87) |
| $R_{pim}$ | 0.035 (0.842) | 0.063 (0.956) | 0.050 (0.617) | 0.057 (0.951) | 0.030 (0.930) |
| CC 1/2 | 0.999 (0.287) | 0.998 (0.320) | 0.997 (0.488) | 0.998 (0.308) | 1.000 (0.343) |
| $I / \sigma I$ | 9.5 (0.4) | 5.9 (0.3) | 8.2 (0.8) | 6.3 (0.3) | 10.1 (0.4) |
| Completeness (%) | 99.2 (96.9) | 100.0 (99.9) | 100.0 (99.7) | 100.0 (98.8) | 99.7 (98.5) |
| Multiplicity | 13.2 (12.6) | 13.1 (12.6) | 13.4 (12.6) | 13.1 (12.0) | 13.2 (13.3) |
| ***Refinement*** | | | | | |
| Resolution (Å) | 35.45 – 0.94 | 36.07 – 1.22 | 35.54 – 1.01 | 36.03 – 1.24 | 41.60 – 0.87 |
| No. reflections | 153778 | 70861 | 126567 | 68096 | 195232 |
| $R_{work}$ / $R_{free}$ | 13.89/15.02 | 16.98/20.88 | 12.95/15.00 | 16.32/19.71 | 13.01/14.11 |
| #. non-H atoms | | | | | |
| Protein | 2106 | 2071 | 2085 | 2081 | 2118 |
| Solvent | 435 | 338 | 355 | 349 | 382 |
| Ligand – NXL | 34 | 17 | 17 | 17 | 17 |
| Ligand – FYG | 17 | 17 | 17 | N/A | N/A |
| B-factors ($Å^2$) | | | | | |
| Protein | 10.7 | 17.0 | 10.5 | 16.8 | 10.6 |
| Solvent | 23.6 | 31.4 | 26.7 | 31.9 | 24.5 |
| Ligand – NXL | 17.4 | 15.9 | 9.71 | 17.5 | 10.5 |
| Ligand – FYG | 18.9 | 22.0 | 14.8 | N/A | N/A |
| R.m.s. deviations | | | | | |
| Bond lengths (Å) | 0.011 | 0.009 | 0.007 | 0.008 | 0.009 |
| Bond angles (°) | 1.299 | 1.097 | 1.062 | 1.040 | 1.055 |
| Ramachandran (%) | | | | | |
| Outliers | 0.39 | 0.39 | 0.39 | 0.39 | 0.39 |
| Favoured | 98.44 | 98.05 | 98.05 | 98.05 | 98.44 |

Values in parentheses are for the high-resolution shell; NXL – covalently reacted avibactam; FYG – intact non-covalently bound avibactam

**Table S5. Room temperature serial synchrotron crystallography data collection and refinement statistics for presoaked CTX-M-15 crystals**

| **Time point** | 600 s |
|---|---|
| **PDB accession** | 9TOH |
| ***Data collection*** | |
| Beamline | DLS I24 |
| Space group | $P2_12_12_1$ |
| Molecules/ASU | 1 |
| Cell dimensions | |
| *a*, *b*, *c* (Å) | 45.27, 45.99, 118.83 |
| α=β=γ (°) | 90 |
| # 'chips' | 1 |
| # integrated lattices | 9888 |
| Wavelength (Å) | 0.9999 |
| Resolution (Å) | 59.414 – 1.70 (1.7293 – 1.70) |
| $R_{split}$ | 14.3 (32.5) |
| CC 1/2 | 97.1 (79.6) |
| $I$ / σ$I$ | 4.881 (2.228) |
| Completeness (%) | 100.0 (99.93) |
| Multiplicity | 43.56 (13.71) |
| ***Refinement*** | |
| Resolution (Å) | 59.41 – 1.70 |
| No. reflections | 28076 |
| $R_{work}$ / $R_{free}$ | 16.48/12.69 |
| # non-H atoms | |
| Protein | 2030 |
| Solvent | 222 |
| Ligand – NXL | 17 |
| Ligand – FYG | 17 |
| B-factors (Å$^2$) | |
| Protein | 16.7 |
| Solvent | 34.3 |
| Ligand – NXL | 11.9 |
| Ligand – FYG | 22.7 |
| R.m.s. deviations | |
| Bond lengths (Å) | 0.009 |
| Bond angles (°) | 1.039 |
| Ramachandran (%) | |
| Outliers | 0.39 |
| Favoured | 98.44 |

Values in parenthesis are for the high-resolution shell; NXL – covalently reacted avibactam; FYG – intact non-covalently bound avibactam

**Table S6. Active site interactions and distances (in Å) in CTX-M-15 crystal structures.**
Schematic below shows overview of active site interactions and atom numbering.

| Atom 1 | Atom 2 | 1.3 s PAL-XFEL | 1.3 s DLS | 10 s cryo conf A | 10 s cryo conf B | 300 s RT | 3 h soak, 10 s wash | 3 h soak, 1800 s wash | 24 h |
|---|---|---|---|---|---|---|---|---|---|
| NXL N6 | Ser130 Oγ | 3.4 | 3.3 | 2.4 | 2.9 | 3.0 | 2.9 | 2.9 | 2.9 |
| NXL O14 | Ser237 Oγ | 3.5 | 3.9 | 3.0 | 3.2 | 2.9 | 3.2 | 2.8 | 3.4 |
| NXL O15 | Thr235 Oγ | 3.0 | 3.0 | 2.6 | 2.6 | 2.7 | 2.7 | 2.6 | 2.6 |
| NXL O16 | Ser130 Oγ | 2.7 | 2.9 | 3.2 | 3.3 | 3.7 | 3.5 | 3.4 | 3.5 |
| NXL O16 | Lys234 Nζ | 3.3 | 3.2 | 3.2 | 3.2 | 3.6 | 3.4 | 3.3 | 3.4 |
| NXL O9 | Ser70 N | 2.7 | 2.7 | 2.6 | 2.7 | 2.7 | 2.6 | 2.7 | 2.7 |
| NXL O9 | Ser237 N | 2.9 | 2.9 | 3.0 | 2.9 | 2.9 | 2.9 | 2.9 | 2.9 |
| NXL O11 | Asn132 Nδ | 2.9 | 2.8 | 2.9 | 2.8 | 2.8 | 2.9 | 2.8 | 2.9 |
| NXL O11 | Asn104 Nδ | 3.2 | 3.9 | 3.0 | 3.2 | 3.2 | 3.0 | 3.0 | 3.0 |
| NXL O11 | Asn170 Nδ | 3.2 | 3.1 | 3.5 | 2.8 | 3.4 | 3.5 | 3.4 | 3.4 |
| NXL N12 | FYG O14 | 3.0 | 3.1 | 3.4 | 3.5 | 3.3 | 3.4 | N/A | N/A |
| NXL N12 | FYG O9 | 3.0 | 2.9 | 2.9 | 3.0 | 3.0 | 3.0 | N/A | N/A |
| FYG O9 | Asn104 Nδ | 3.1 | 2.9 | 3.1 | 3.1 | 3.1 | 3.1 | N/A | N/A |
| Ser130 Oγ | Lys234 NZ | 2.9 | 3.0 | 2.8 | 2.8 | 2.8 | 2.8 | 2.8 | 2.8 |
| Ser130 Oγ | Lys73 NZ | 3.1 | 3.0 | 2.9 | 2.9 | 2.8 | 2.8 | 2.8 | 2.8 |

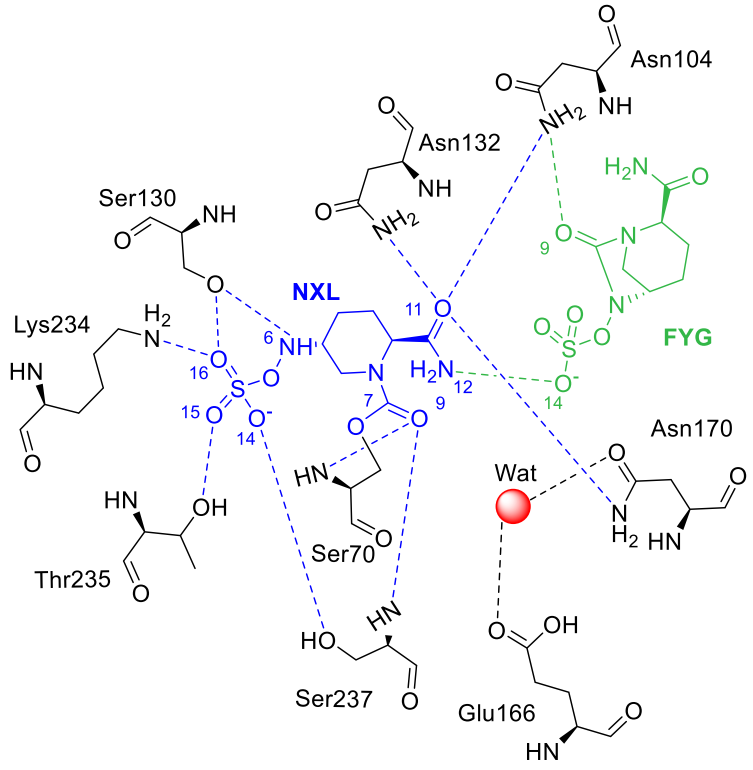

**Table S7. Key distances and angles in X-ray structures optimised with Quantum Mechanics/Molecular Mechanics at the Density Functional Theory level.**

| System | O9-C7-N6 Angle (°) | H-C5-N6-H Dihedral (°) | C7-N6 Distance (Å) | Lys73$^{N\zeta}$ – Ser130$^{H\gamma}$ Distance (Å) | Ser130 – Avi$^{N6-H}$ Distance (Å) | Glu166$^{O\varepsilon2}$ -DW$^{O}$ Distance (Å) | DW$^{O}$-C7 Distance (Å) |
|---|---|---|---|---|---|---|---|
| **1.3 s (PAL-XFEL)** | 124.2 | 64.2 | 3.9 | 1.7 | 4.4 | 2.6 | 3.2 |
| **10 s conf. A (100 K)** | 117.9 | 280.5 | 3.4 | 1.7 | 1.9 | 2.6 | 3.3 |
| **10 s conf. B (100 K)** | 110.5 | 306.2 | 3.0 | 1.7 | 1.9 | 2.6 | 3.2 |
| **600 s (room temp.)** | 108.6 | 301.4 | 2.9 | 1.8 | 1.9 | 2.6 | 3.2 |
| **24 Hours (100 K)** | 112.5 | 309.4 | 2.9 | 1.7 | 1.8 | 2.6 | 3.2 |
